# Understanding the structure and mechanics of the sheep calcaneal enthesis: a relevant animal model for tissue engineering applications

**DOI:** 10.1101/2024.12.24.630234

**Authors:** Alberto Sensini, Luca Raimondi, Albano Malerba, Carlos Peniche Silva, Andrea Zucchelli, Alexandra Tits, Davide Ruffoni, Stéphane Blouin, Markus A. Hartmann, Martijn van Griensven, Lorenzo Moroni

**Affiliations:** Department of Complex Tissue Regeneration (CTR), MERLN Institute for Technology-Inspired Regenerative Medicine, Maastricht University P.O. Box 616, Maastricht 6200 MD, The Netherlands; Department of Cell Biology-Inspired Tissue Engineering (cBITE), MERLN Institute for Technology-Inspired Regenerative Medicine, Maastricht University, P.O. Box 616, 6200 MD Maastricht, The Netherlands; Department of Industrial Engineering, Alma Mater Studiorum - Università di Bologna, Bologna, Italy; Advanced Mechanics and Materials – Interdepartmental Center for Industrial Research (CIRI-MAM), Alma Mater Studiorum—Università di Bologna, Bologna, Italy; Mechanics of Biological and Bioinspired Materials Laboratory, Department of Aerospace and Mechanical Engineering, University of Liège, Quartier Polytech 1, Allée de la Découverte 9, 4000, Liège, Belgium; Ludwig Boltzmann Institute of Osteology, Hanusch Hospital of OEGK and AUVA Trauma Centre Meidling, 1st Medical Department Hanusch Hospital, Vienna, Austria

**Keywords:** Tendon tissue, enthesis tissue, Digital Image Correlation, Mechanical Tensile and Cyclic Tests, Nanoindentation, Scanning Electron Microscopy, Histology

## Abstract

Tendon/enthesis injuries are a worldwide clinical problem. Along the enthesis, collagen fibrils show a progressive loss of anisotropy and an increase in mineralization reaching the bone. This causes gradients of mechanical properties. The design of scaffolds to regenerate these load-bearing tissues requires of being validated *in vivo* in relevant large animal models. The sheep tendon of triceps surae muscle is an optimal animal model for this scope with limited knowledge about its structure and mechanics. We decided to understand in-depth its structure and full-field mechanics. Collagen fibrils morphology was investigated via scanning electron microscopy revealing a marked change in orientation/dimensions passing from tendon to enthesis. Backscatter electron images and nanoindentation at the enthesis/bone marked small gradients of mineralization at the mineralized fibrocartilage reaching 27%wt and indentation modulus around 17-30 GPa. The trabecular bone instead had indentation modulus around 15-22 GPa. Mechanical tensile tests with digital image correlation confirmed the typical non-linear behavior of tendons (failure strain = 8.2±1.0%; failure force = 1369±187 N) with maximum principal strains reaching mean values of ε_p1_∼7%. The typical auxetic behavior of tendon was highlighted by the minimum principal strains (ε_p2_∼5%), progressively dampened at the enthesis. Histology revealed that this behavior was caused by a local thickening of the epitenon. Cyclic tests showed a force loss of 21±7 % at the last cycle. These findings will be fundamental for biofabrication and clinicians interested in designing the new generation of scaffolds for enthesis regeneration.

**Graphical Abstract:** 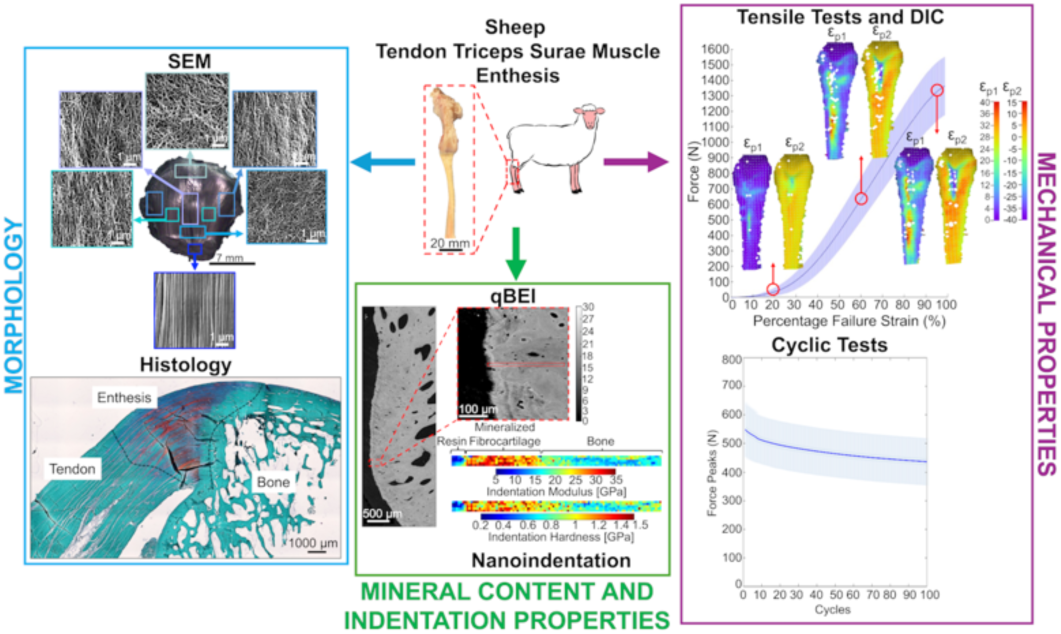

**Statement of Significance:** Tendon and enthesis lesions are a clinical problem. To validate scaffolds for these applications large animal models are needed. Sheep tendon of triceps surae muscle is an optimal site for this scope. However, little is known about its extracellular matrix structure and mechanical properties. This work investigates the structure and mechanics of this tissue from different points of view. Scanning electron microscopy and histology studied its extracellular matrix morphology and composition. Backscattered electron images and nanoindentation assessed gradients of mineralization and stiffness at the enthesis. Mechanical tensile and cyclic tests coupled with digital image correlation elucidated its mechanics and superficial strain distribution. These findings will be fundamental for biofabrication and clinician experts to design innovative scaffolds to regenerate the enthesis.

## 1. Introduction

The regeneration/replacement of injured tendons and ligaments (T/L) is an extremely relevant societal need worldwide, with 30 million cases estimated annually. The healthcare expenditure to manage these injuries exceeds €115 billion in Europe and $30 billion in the USA per year [1]. Among these, a relevant site of injury is the T/L-to-bone insertion (enthesis), often related to rotator cuff diseases, tennis elbow, jumper’s knee, and Achilles tendinosis [2,3]. Enthesis injuries often result in acute disability and may predispose joints to osteoarthrosis, a disease estimated to affect over 70% of people aged 55–78 [4,5]. The failure rates after surgeries for enthesis repair are extremely high, ranging from 10% up to 94% depending on the anatomical site (e.g., rotator cuff: 20-94%) [6]. The reason behind these criticalities resides in the complex hierarchical organization of the extracellular matrix (ECM) of the T/L, fibrocartilage and bone tissues at the enthesis. Focusing on tendons, they are mainly composed of highly packed and axially oriented collagen type I fibrils grouped in more complex structures such as fibers, fascicles, and tertiary fiber bundles, that are protected by collagen membranes (i.e., endotenon and epitenon) [7,8]. The enthesis has an additional complexity as it features multiple tissues confined in a very small region. From a biomechanical viewpoint, one important task of the enthesis is to reduce stress concentration, a well-known problem of bi-material junctions in engineering. This is achieved by compositional and microstructural modifications, possibly resulting into a gradual transition of local mechanical properties (e.g. stiffness) passing from the tendon to the bone tissue. In light of this, the enthesis is composed of different regions where a progressive loss of anisotropy of the collagen fibrils, from the tendon to the bone, is matched with a gradient of mineralization [9–11]. Specifically, the four different regions are the tendon tissue, the non-mineralized fibrocartilage (n-mFC), the mineralized fibrocartilage (mFC) and the bone tissue (Fig. 1A) [12]. Focusing on the ECM structure and composition, the first region consists of axially aligned collagen type I fibrils [9]. The n-mFC shows a relevant amount of collagen types II and III with small amounts of collagen types I and X, decorin, and aggrecan [13,14]. In terms of morphology, n-mFC assumes a typical conical structure [15], functional to reduce the interfacial stress concentrations, while collagen fibrils progressively increase their randomicity. In the mFC instead, collagen type II, X and aggrecan are the most prevalent [13,14] and fibrils further increase their randomic orientation. In the bone tissue instead, the ECM is mainly composed of mineralized collagen type I reinforced with nanometer-sized mineral crystals [12] with the typical trabecular and cortical organization.

**Fig. 1.**
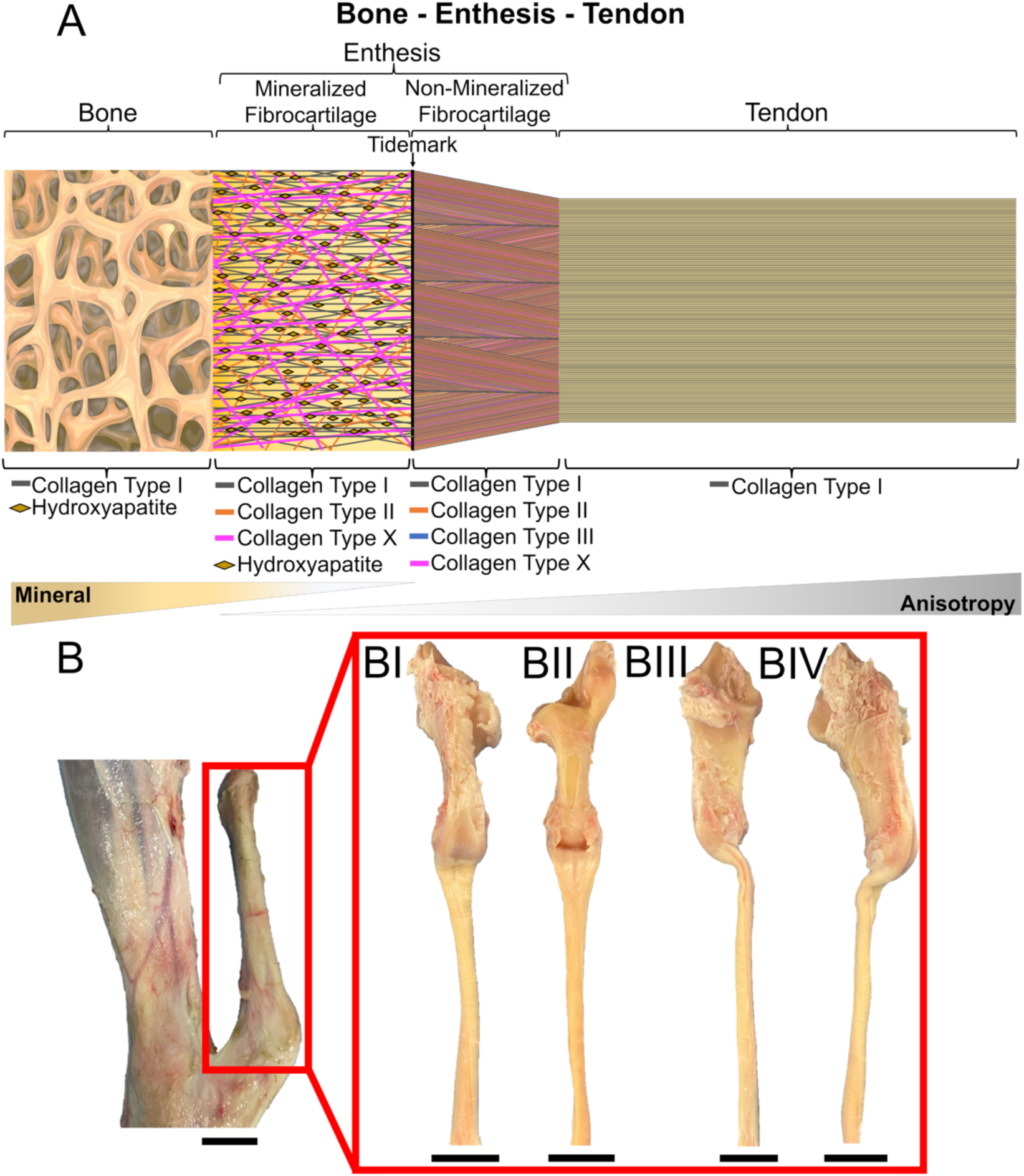
Sheep TTSM and calcaneal bone. A) Organization, composition and anisotropy/mineral gradients of ECM; B) Image of a sheep right posterior leg with its calcaneus and Achilles tendon (scale bar = 20 mm). BI-BIV) Different views of a sample after preparation, consisting of the calcaneal bone with the TTSM attached. From each specimen, the bursa, ligaments, and surrounding membranes were removed (scale bar length =20 mm).

This sophisticated structural/mineral architecture of the ECM at the enthesis is mirrored into gradients of mechanical properties. In the literature, enthesis were investigated at different length scales with several techniques [16], such as tensile/fatigue tests [17–19] and nanoindentation [11]. the relationship between composition, fiber arrangement and mechanical gradients, in both animal and human cadaveric samples, mechanical tensile tests were coupled with full-field contactless imaging techniques, such as Digital Image Correlation (DIC) [20–23], Digital Volume Correlation (DVC) [24,25] and confocal microscopy [15]. Interestingly, DIC and DVC were also able to detect gradients of auxetic behavior at the tendon and the enthesis sides both in human Achilles tendons [23] and in mice [24]. Considering the complex interplay between multiscale morphology, collagen fibrils composition/architecture and the related complex mechanics of the enthesis, its surgical intervention often leads to unsatisfactory outcomes. For this reason, researchers are constantly working on the design of innovative biomimetic scaffolds to regenerate these injured tissues [26–28]. Despite promising *in vitro* results, scaffolds must be validated through *in vivo* implantation in relevant animal models, before reaching clinical trials in humans*. In vivo* studies to test tendon and enthesis scaffolds often involve small animals such as rabbits, mice and rats or, when particularly promising, large animals such as dogs and mostly sheep [29–31]. The typical anatomical sites of implantation in both small/large animal studies are the rotator cuff, the patellar and the Achilles tendon [29–32]. Specifically, the Achilles tendon of sheep/goats has been demonstrated to be the most relevant model to study the tissue regeneration mechanisms and scaffolds performances being its collagen composition and weight-bearing capacity similar to the human one [33–37]. The posterior Achilles tendon of the sheep has a complex structure since it is composed by two major tendons, the superficial digital flexor tendon (SDFT) and the tendon of the triceps surae muscle (also called calcaneal tendon or TTSM), grouped together by a peritenon tissue membrane [38]. While the SDFT overpasses the calcaneal bone, producing an enthesis at the foot, the TTSM attached to the calcaneal bone through a marked enthesis [38]. This makes TTSM a suitable anatomical site to recapitulate the human Achilles tendon and its enthesis at the calcaneus bone.. Although some studies focused on the whole sheep Achilles tendon to study the tissue regeneration performances of commercial grafts and surgical sutures [35,36], the fibrillar organization, mineral enthesis gradients and superficial local strain mechanics of the tendon-enthesis-bone complex of the TTSM (i.e. the most relevant tendon to mimic the human Achilles one), remain practically unexplored so far.

Aiming at further understanding the morphology of the sheep TTSM and its enthesis, a panel of morphological and mechanical investigations were performed. At first a Scanning Electron Microscopy (SEM) and histological investigations were performed to better understand the ECM composition and organization of TTSM and its enthesis. The mineral content and mechanical properties at the mFC and bone side of enthesis were evaluated via quantitative backscattered electron imaging (qBEI) and nanoindentation. Finally, the mechanical properties of the calcaneal tendon were characterized by a number of distinct experiments: mechanical tensile test to failure coupled with a DIC investigation of the tendon-enthesis-bone complex, to understand the overall mechanical response and full-field superficial strain distribution of the tissues involved; cyclic tests were performed to detect the loss of force of the calcaneal tendon-enthesis complex.

## 2. Materials and methods

### 2.1. Animal samples collection and preparation

20 right posterior legs were extracted from sheep (mean age 6 years old and 50 kg weight) sacrificed for alimentary purposes. Legs were cut from the distal side of the posterior knee up to the proximal side of the femur as shown in Fig. 1B. Skin, muscles, surrounding bones/tissues, peritenon, and surrounding ligaments were removed to insulate the TTSM tendon-enthesis-bone complex (schematic of the ECM organization/composition in Fig. 1A). Then, the SDFT, endotenon, and bursa were also removed, leaving the calcaneal bone, enthesis, and tendon only (Fig. 1BI - BIV). Specimens were stored frozen at -30°C and then defrosted in saline solution before being prepared for the different tests.

### 2.2. SEM investigation

To study the ECM organization of TTSM, a SEM investigation was performed. At first, to investigate the surface of the TTSM at the enthesis region, one sample was cut at the bone side, leaving approximately 5 mm of tendon attached to the bone (Fig. 3A). Concurrently, to analyze the inner section of TTSM, a second sample was cut longitudinally to make its cross-section visible (Fig. 4). Samples were fixed and decellularized adopting a consolidated procedure [39]. In short, samples were fixed in 4% glutaraldehyde (Merck) in 0.1M phosphate buffer (Merck) for 2 days inside a 4°C fridge. Then, samples were rinsed in distilled water and incubated in 5% NaOH solution at room temperature for 5 days. After an additional rinsed distilled water, samples were immersed in 1% tannic acid for 4 hours at room temperature and finally rinsed again in distilled water overnight. After three immersions in 0.1M sodium cacodylate buffer, samples were immersed in 1% osmium tetroxide in 0.1M sodium cacodylate buffer for 1 hour. After three additional rinses in 0.1M sodium cacodylate buffer, samples were immersed in different ethanol concentrations (70%, 90%, 100%). Finally, two immersions of 30 minutes in hexamethyldisilazane (Merck) were performed, before coating the samples with carbon and mounting the specimens in stubs with silver glue. TTSM samples were investigated by using a SEM (JEOL JSM-IT200, Peabody, MA, USA) operated at 10 kV. To study the diameter distribution and the orientation of the ECM fibrils in six different regions of TTSM enthesis surface were defined: the tendon region (RI), the region below the enthesis (RII), the center of enthesis (RIII), the body of enthesis (RIV), the sides of enthesis (RV) and the bone region (RVI). Then, 5 images, for each of the 6 regions of samples detected (Fig. 3), were acquired (magnification: 10000x). To study the orientation of fibrils in the different regions of TTSM samples, the Directionality plugin of ImageJ was applied (using the Local Gradient Orientation method of the plugin), adapting a consolidated procedure [40]. The diameter distribution per each region of interest was obtained by calculating 200 fibrils’ diameters. The measurement of the thickness of the fibrocartilage fibrils layer covering the TTSM fascicles at the enthesis level was calculated by using ImageJ as the mean ± SD of 20 measurements.

### 2.3. Histology

Tendon-to-bone units (n = 2) were fixed for 48h using 4% paraformaldehyde (SigmaAldrich). Consequently, samples were rinsed with PBS (Thermo Fisher Scientific) and decalcified for 35 days in 10% buffered EDTA solution at 4 degrees with buffer exchanged every 2 to 3 days. Decalcified samples were dehydrated in ethanol series (50%, 70%, 96%, and 100%) and embedded in parafine. Then, samples were longitudinally sectioned at a thickness of 7 µm using a Leica RM2165 microtome (Leica Biosystems) and stored at 4 degrees. Safranin-O staining was performed on samples following the standard protocol indicated by the manufacturer and a previous consolidated method [32]. Briefly, samples were deparaffinized in NeoClear-xylene substitute and rehydrated in descending ethanol series (100%, 96%, 70% and 50%) and distilled water. Samples were stained with hematoxylin solution for 10 min, followed by 10 min wash in running tap water and a 5 min staining with 0.1% fast green solution (Sigma-Aldrich). Later, samples were rinsed with 0.1% acetic acid and stained with 0.1% safranin O solution for 10 min (Sigma-Aldrich). Thereafter samples were dehydrated in ascending ethanol series (70%, 95%, and 100%), cleared using NeoClear-xylene substitute, and mounted using UltraKit mounting media. Stained samples were imaged with an inverted Nikon Ti-S/L100 microscope.

### 2.4. Mineral content and local tissue mechanical properties

#### 2.4.1. Samples preparation

Right posterior sheep calcaneal bone and TTSM enthesis (n = 3) were prepared to expose representative sagittal cross-sections of the tendon-bone insertion. Samples were trimmed using a diamond saw (Buehler Isomet) to fit the embedding molds (30 mm diameter). A 10-day dehydration protocol was followed, involving immersion in progressively concentrated ethanol and acetone solutions. For embedding, samples were immersed in MMA for 2 days, infiltrated with liquid PMMA resin in a refrigerator for another 2 days, and then allowed to harden in an oven for 6 days, with temperatures gradually increasing from 34°C to 50°C. After cutting the samples to expose the enthesis, they were ground with P1500 and P2500 grit grinding papers, then polished (Buehler MetaServ 250) in three steps using diamond grains of 3 μm, 1 μm, and 0.25 μm, under glycol lubrication. Finally, the samples were cleaned with petroleum ether.

#### 2.4.2. qBEI analysis

To measure the mineral content, we performed a qBEI analysis. This is a well-established technique based on SEM, measuring the intensity of the backscattered signal, which primarily reflects the local calcium concentration [41,42]. We used a Field Emission Scanning Electron Microscope (FESEM, Supra40, Zeiss) at 20 kV with a working distance of 10 mm and a scan speed of 90 seconds per frame. Images were captured at 130x magnification, producing a pixel size of 0.88 μm within a 1024×768-pixel window. To quantitatively assess calcium content, the FESEM was calibrated with carbon and aluminum standards using a previously established method [42]. The reference materials were assigned gray values of 25 and 225, corresponding to osteoid (0 wt% Ca) and pure hydroxyapatite (39.86 wt% Ca), respectively. This calibration allowed the conversion of gray level (GL) images into spatial maps of mineral content, expressed as calcium weight percentage, using the equation:

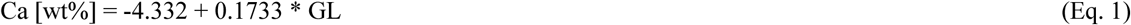

We excluded signals from the embedding resin and non-mineralized tissues by setting a minimum mineral content threshold of 5.2 wt%.

#### 2.4.3. Nanoindentation

After the qBEI analysis, samples were characterized by nanoindentation (Triboindenter TI 950, Bruker) to obtain local mechanical properties across the enthesis expressed as indentation modulus and hardness [43]. A Berkovich diamond probe and a displacement-controlled trapezoidal load function (8 - 12 - 8 s) with a maximum displacement of 200 nm were used to probe the tissues [11]. Several indentation grids were performed across the bone-mFC interface for a total of 1250 indents, with regions selected based on the qBEI maps and optical microscopy. The spacing between adjacent indents ranged from 2 µm to 3 µm. The probe contact area was calibrated using fused quartz, and the load-depth curves were analyzed using the Oliver-Pharr method [43] to determine indentation modulus (E_r_) and hardness (H). The nanoindentation results were processed using a custom MATLAB script to obtain 2D maps of the mechanical properties. These maps were interpreted with the help of region-matched qBEI maps, allowing the classification of indents falling into bone and mFC, as well as discarding indents located in cracks or pores. Finally, nanoindentation was also performed in trabecular bone to have additional values away from the enthesis. A single grid of 5 x 5 indents with a spacing of 4 μm both vertically and horizontally, resulting in a total dimension of 20 x 20 μm, has been performed for each sample (n=3 trabeculae). The post-processing of the data was similar as previously described.

### 2.5. Tensile and cyclic mechanical tests

#### 2.5.1 Samples preparation for mechanical tensile and cyclic tests

A custom-made setup (Fig. 2) has been specifically designed to secure the bony side of samples and guarantee the physiological alignment of each specimen along the loading axis of the testing machine (with an angle of 30° between TTSM and the calcaneal bone [38] as shown in Fig. 2A). Before the test, the bony side of the specimen was immersed in polymethylmethacrylate (PMMA) directly inside the custom-made setup. To prevent slippage, the proximal sides of tendons (TTSM side) were fixed in 25 mm of emery cloth with cyanoacrylate glue and stapler pins, and finally clamped in a standard tensile clamp for mechanical tests.

**Fig. 2.**
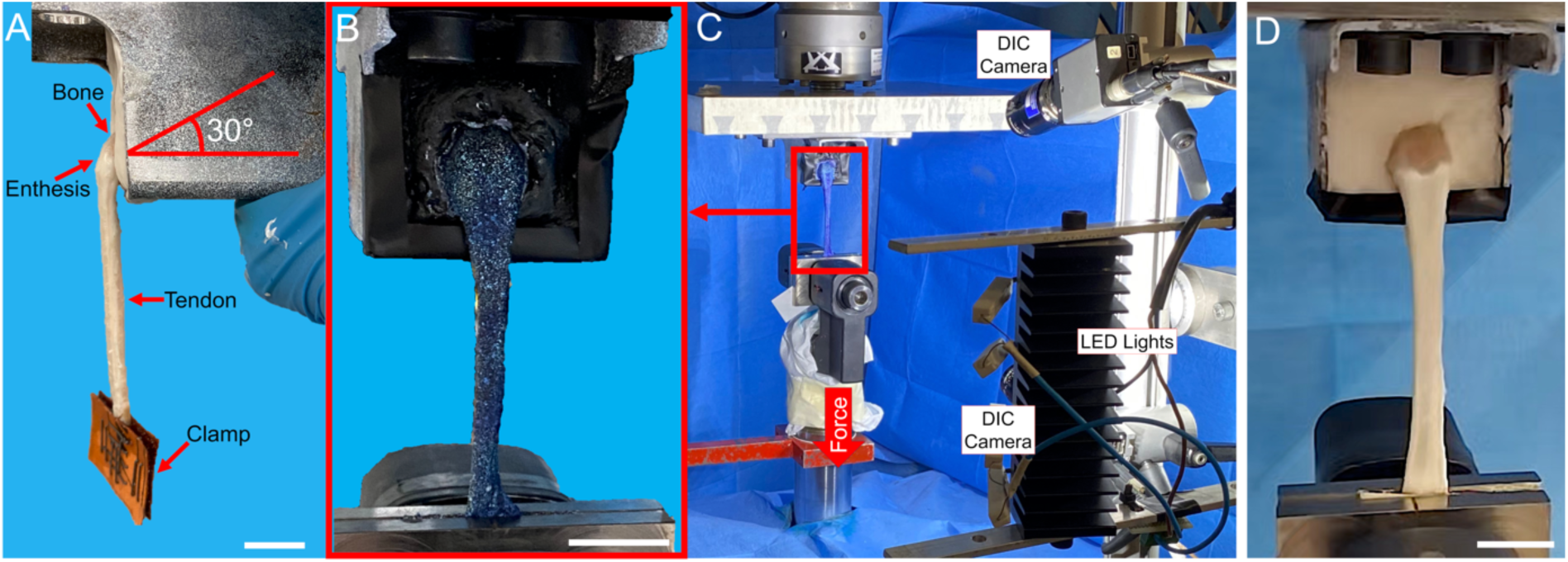
Setup for the mechanical tests and DIC. A) Mounting setup of samples. B) Zoom-in at a tendon of triceps surae muscle (TTSM) sample, stained with methylene blue and with a white speckle pattern, mounted in the custom setup for the mechanical DIC test. C) Overview of the DIC system and the testing setup for tensile tests. D) TTSM sample mounted in the testing machine for the cyclic tests (scale bar = 20 mm for all images).

The gauge length of each specimen was calculated as mean and standard deviation (SD) between 3 measurements from the calcaneal tuberosity (in correspondence of the enthesis) up to the clamp. The specimens’ cross-section was assumed rectangular and was calculated as mean ± SD between 30 measurements from high-resolution pictures of each side of the specimens. The gauge length resulted in a mean ± SD between all samples of 74.2 ± 8.01 mm, measured from the calcaneal tuberosity (i.e. enthesis) to the lower clamp.

#### 2.5.2. Mechanical tensile tests and DIC

Samples (n = 6) were monotonically loaded to failure into a servo-hydraulic testing machine (8032, Instron, UK) equipped with a 10 kN load cell (Instron, UK), operated in displacement control. The crosshead speed was adjusted to ensure a strain rate of 1% s^-1^ for each specimen. Before testing, samples were pre-conditioned with five cycles at 2% strain to allow the adjustment of collagen fibrils. Force and displacement were recorded at a sampling rate of 1 kHz. The nominal strain (i.e. global strain) of each sample was calculated as:

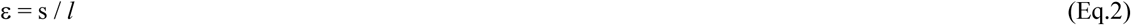

Where *s* is the relative displacement of the crosshead of the testing machine and *l* is the gauge length of the specimen of interest.

To calculate the stress (σ), the cross-sectional area of each TTSM was approximated as rectangular. For each specimen, 20 measurements of the base (b = width) and the high (h = thickness) were calculated by using ImageJ on 4 high-resolution images along the gauge length of each specimen (see Fig. 1BI-BIV) for the views of a typical specimen). The mean ± SD of the cross-sectional area between all samples resulted to be 44.5 ± 10.3 mm^2^.

Stress was finally calculated as follows:

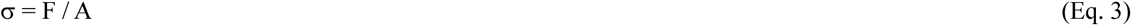

Where *F* was the force and *A* the cross-sectional area (defined as A = b x h) of each sample. The following mechanical properties were obtained: elastic modulus (E), stiffness (K), force at failure (F_F_), stress at failure (σ_F_), strain at failure (ε_F_), energy at failure (J_F_) and unit work to failure (W_F_). In particular, J_F_ and W_F_ were calculated, with the trapezoidal rule, as the area under the force-displacement and stress-strain curves respectively.

To identify the point where the first groups of fibrils of TTSM started damaging macroscopically, the inflection point was calculated by applying a previously consolidated method [44]. In brief, this value is defined as the point inside the linear region of the stress-strain curve of TTSM, where the tissue passes from strain-stiffening to strain-softening. At first, by using a caps-based MATLAB script, the mechanical data were fitted with a cubic smoothing spline. The smoothing parameter of *csaps* function of MATLAB was set to 0.7, guaranteeing a smooth first and second derivative of curves. The inflection point was computed by identifying the zero point of the second derivative. Mean ± SD of inflection point strain (IPε), force (IPF), stress (IPσ) and percentage of failure strain (IP_%εF_) were calculated.

To investigate the distribution of the local strains on the surface of samples during the tensile test, a parallel DIC investigation was carried out by measuring the maximum (ε_p1_) and minimum (ε_p2_) principal strain. Tests were monitored by 2 cameras (5 MegaPixels, 2440 × 2050 pixels, 8-bit) (Fig. 2B) of a commercial 3D-DIC system (Q400, Dantec Dynamics, Denmark) equipped with high-quality metrology-standard 35 mm lens (Apo-Xenoplan 1.8/35, Schneider−Kreuznach, Germany; 135 mm equivalent) for a stereoscopic view. To ensure effective illumination of samples, a custom-designed array of LEDs (10,000 lumens in total) was used. To mitigate correlation artifacts associated with metallic components and white polymethylmethacrylate in the fixture, black insulating tape and paint were utilized. Before each session, the system was calibrated with a proprietary calibration target (Mod. Al4-BMB-4×4, Dantec Dynamics), achieving 3D-residuum values from 0.050 to 0.150. The process involves capturing a sequence of images of the subject from various angles to determine the physical size of the measuring area. This is followed by correcting any distortions caused by the lens and compensating for parallax effects. Care was taken in optimizing the system to achieve the best compromise between accuracy and spatial resolution. For DIC measurements, to ensure a dark background while avoiding cracks in the painting for the large deformations applied, samples were previously painted with methylene blue solution (4 g of methylene-blue per 100 ml of water) following the protocol reported by Lionello et al. [45]. Then, a white speckle pattern was applied by spaying with an airbrush-airgun (AZ3 HTE 2, nozzle 1.8 mm, Antes Iwata, Italy), a water-based paint (Q250201 Bianco Opaco, Chrèon, Italy) diluted at 40% with water (Fig. 2A). Care was taken in optimizing the pattern to have a speckle size of approximately 3-5 pixels with a pattern density of about 50% to provide low noise-affected measurements [46]. Before testing, the specimens were subjected to hydration using saline solution, meticulously applied via a spray bottle. This ensured the integrity of the speckle pattern and the hydration of samples, fundamental for the correlation accuracy of the DIC system.

The samples were positioned 500 mm away from the cameras, which were set up vertically to ensure that both the tendon and enthesis could be captured during the test. The correct field of view was adjusted to 105 mm by 125 mm, resulting in a pixel size of 50 µm. This field of view was wide enough to encompass the entire sample during the test. Images were captured at a frame rate of 10 Hz. To assess the measurement uncertainties, a zero-strain analysis was performed by acquiring two images of the unloaded samples for each specimen and analyzed. The Kolmogorov-Smirnov test was used to check that the errors followed a Gaussian distribution. The systematic and random errors were calculated by computing the mean and SD of ε_p1_ and ε_p2_ in the zero-strain configuration for each sample [47–49].

Images were post-processed by ISTRA 4D software (v.4.3.1, Dantec Dynamics, Denmark) by using a facet size of 21 pixels and a grid spacing of 11 pixels for the highest possible spatial resolution.

To provide a reliable and comparable measurement of the strain of the different samples under investigation, both maximum and minimum principal strains ε_p1_ and ε_p2_ were normalized on ε_F_ (i.e. measured strain at failure of the specimen of interest), obtaining maximum normalized (ε_p1n_) and minimum normalized (ε_p2n_) principal strains as follows:

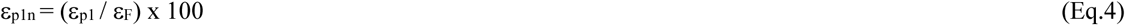

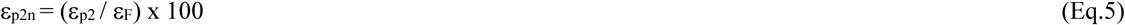

Virtual strain gauges were used to determine averaged values of ε_p1_, ε_p2_ (as well as ε_p1n_, ε_p2n_, and their corresponding maximum and minimum data) to improve the accuracy of the measurements at tailored and distinct regions by directly drawing them on the existing correlated datasets. Three virtual strain gauges (i.e. VSGI-III) were directly drawn on the correlated images following the contours distinctive patterns characterized by RI-RVI. Specifically, VSGI was used in RI to contour all the tendon region, VSGII delimited the enthesis (RII-RV regions), while RVI, the bone region, was delimited inside VSGIII (as defined in section 2.2) (see Fig. 6G). This to have enough correlated points to allow the calculation of averaged ε_p1_ and ε_p2_ values. In fact, working with biological soft tissues in a large-strain regime, the mutual interaction between the stretching of the speckle pattern, the pullout of water from the specimens during the test, and their reduced dimensions facilitated locally the progressive loss of correlation in some points.

#### 2.5.3. Cyclic tests

Samples (n = 6) were cyclically loaded (up to 100 cycles) (Fig. 2C) in displacement control by using a frequency of 1 Hz at 5% nominal strain using the same setup described in section 2.4.2. This strain value was chosen because it is often used to cyclically load scaffolds for tendon tissue regeneration [50,51] and after verifying that this strain value was below the average inflection point identified during the tensile tests (see Table 1). Before the test, a pre-conditioning with five cycles at 2% strain allowed the adjustment of collagen fibrils. To evaluate the progressive loss of force, each force peak (F_P_) at 5% strain was registered per each cycle. The values of single samples and mean ± SD of: the force peaks (F_P_) values, the percentage of the force loss between the first cycle’s peaks of the first packages compared with the first cycle’s peak of each packages (F_Peak_loss_) and the percentage of force peak loss (F_Packages_loss_) between the last cycle’s peaks of two consecutive packages (10 cycles) of the cyclic tests of TTSM samples were also calculated.

**Table 1.** Mechanical tensile properties of TTSM.

| Sample | $F_F$<br>(N) | $\sigma_F$<br>(MPa) | $\varepsilon_F$<br>(%) | E<br>(MPa) | K<br>(N/mm) | $W_F$<br>(J/mm<sup>3</sup>) | IFF<br>(N) | IFσ<br>(MPa) | IFε<br>(%) | IF$\varepsilon_F$<br>(%) |
| --- | --- | --- | --- | --- | --- | --- | --- | --- | --- | --- |
| (n = 6) | 1369±187 | 33±11 | 9.0±1.2 | 531±112 | 309±38 | 0.0012±0.0006 | 84±18 | 28±14 | 7.8±1.8 | 86±11 |

## 3. Results

### 3.1. SEM investigation

The SEM investigation (Fig. 3 and Fig. 4) revealed a collagen fibril orientation and diameter distribution strongly dependent on the specific zone of TTSM, transitioning from the tendon tissue (i.e. RI in Fig. 3) to the other regions of the enthesis (i.e. RII-RV in Fig. 3) and finally up to the bone tissue (i.e. RVI in Fig. 3). Specifically, a prevalent axial alignment of the collagen fibrils in correspondence with the TTSM tendon body (RI in Fig. 3G-3J) was detected, with a frequency of fibrils oriented in the range 0° - 18° = 76 ± 6 % (range 78° - 90° = 1.6 ± 0.3 %) (Fig. 3J) and a mean fibrils diameter of 170 ± 31 nm (Fig. 3G-3I and Fig. 3K). Approaching the TTSM enthesis, the fibril diameter distribution decreased compared with the ones in the tendon tissue (RI) in a region-specific manner (fibrils diameter: RII = 71 ± 17 nm; RIII = 87 ± 23 nm; RIV = 93 ± 25 nm; RV = 97 ± 25 nm; RVI = 91 ± 24 nm). A similar trend was assumed by the superficial orientation of fibrils that markedly changed in the different regions becoming progressively more randomly oriented approaching the enthesis and the bone regions. In RII and RVI collagen fibrils assumed a mostly randomly arranged orientation but with a slightly axial orientation (RII: range 0° - 18° = 31 ± 2 %, range 78° - 90° = 12 ± 1 %; RVI: range 0° - 18° = 30 ± 2 %, range 78° - 90° = 13 ± 1 %) (Fig. 3F, 3C). In RIII and RIV, fibrils assumed a higher axial orientation (RIII: range 0° - 18° = 38 ± 4 %, range 78° - 90° = 10 ± 2 %; RIV: range 0° - 18° = 38 ± 3 %, range 78° - 90° = 9 ± 1 %) (Fig. 3B, 3E) that markedly increased along the points of attachment of TTMS with the proximal and medial sides of the calcaneal bone (RV: range 0° - 18° = 50 ± 3 %, range 78° - 90° = 5.5 ± 0.8) (Fig. 3D).

**Fig. 3.**
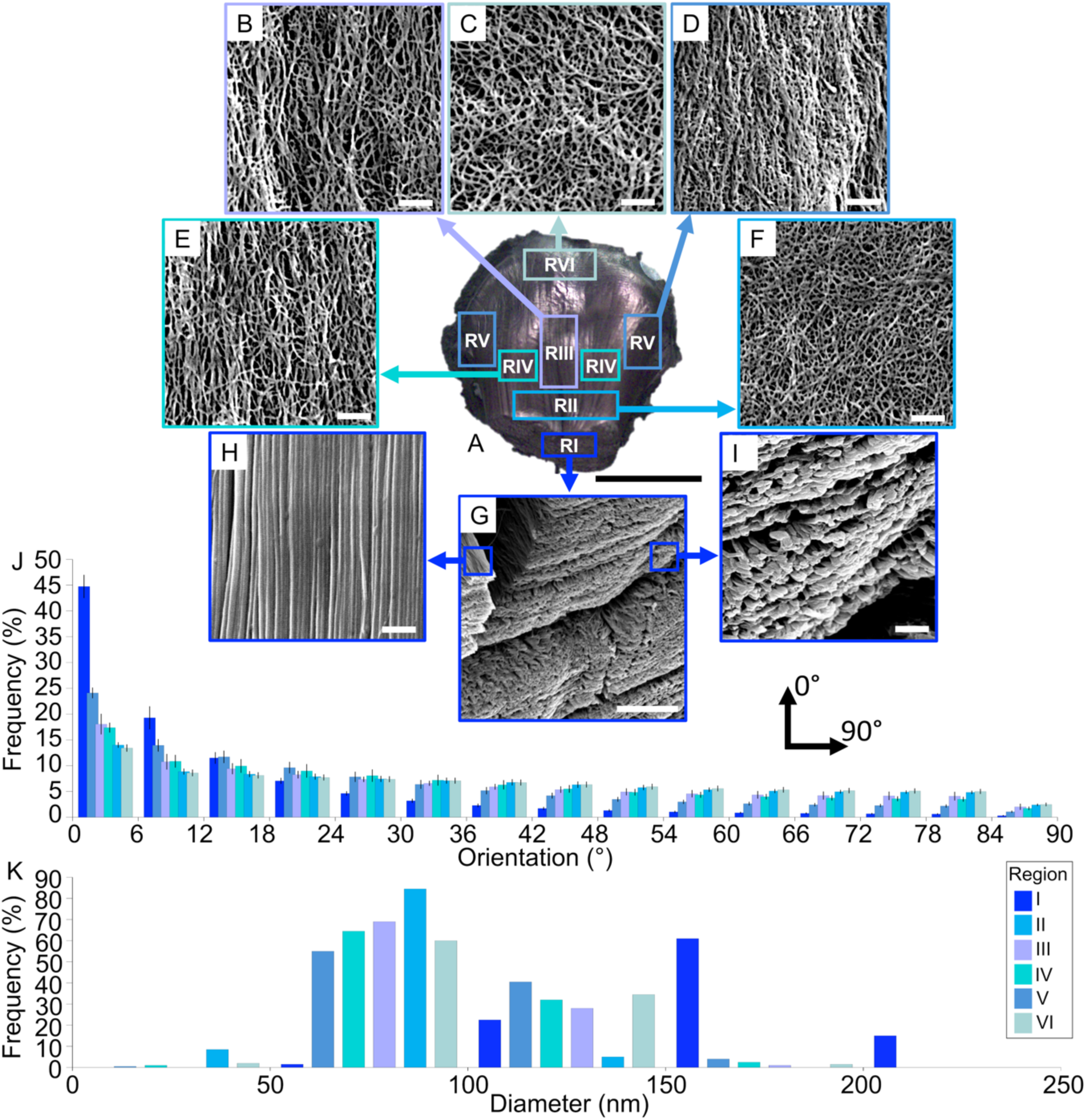
Morphology of the surface of enthesis over the calcaneal bone of TTSM. A) overview of TTSM at the calcaneal bone (scale bar = 7 mm). SEM images of the different regions (RI-RVI) of TTSM enthesis: B) RIII; C) RVI; D) RII; E) RIV; F) RV; G) RI cross-section (scale bar = 10 μm; magnification 2000x); H) RI zoom-in at fibrils surface; I) RI zoom-in at fibrils cross-section (images B-F and H,I scale bar = 1 μm; magnification 10000x). J) Directionality histograms show the distribution of the nanofibers in the five regions of the enthesis of TTSM. An angle of 0° represents collagen fibrils axially aligned with the axis of TTSM, while an angle of 90° represents fibrils perpendicularly oriented with the axis of TTSM. K) Collagen fibrils diameter distribution in the different regions of the enthesis of TTSM.

Looking at the cross-section of TTSM (Fig. 4A), it was possible to see the tendon fascicles (red square) attaching to the bone tissue (red triangle) (Fig. 4B). Moreover, it was possible to see also the typical conical structure of tendon fascicles of enthesis at the n-mFC level (red star) and their collagen fibrils, entering inside the bone tissue with the mFC (red triangle) (Fig. 4C, Fig. 4D). Interestingly, a thick layer of packed random fibrils (thickness ∼ 100 μm) was detected on the surface of TTSM (Fig. 3), extended up to the calcaneal bone (red circles) (Fig. 4E, 4F). Probably this fibrous layer represents a thickening of the epitenon sheath developed to protect the tendon fascicles below from the high friction produced by the interaction with the SDFT in this area.

**Fig. 4.**
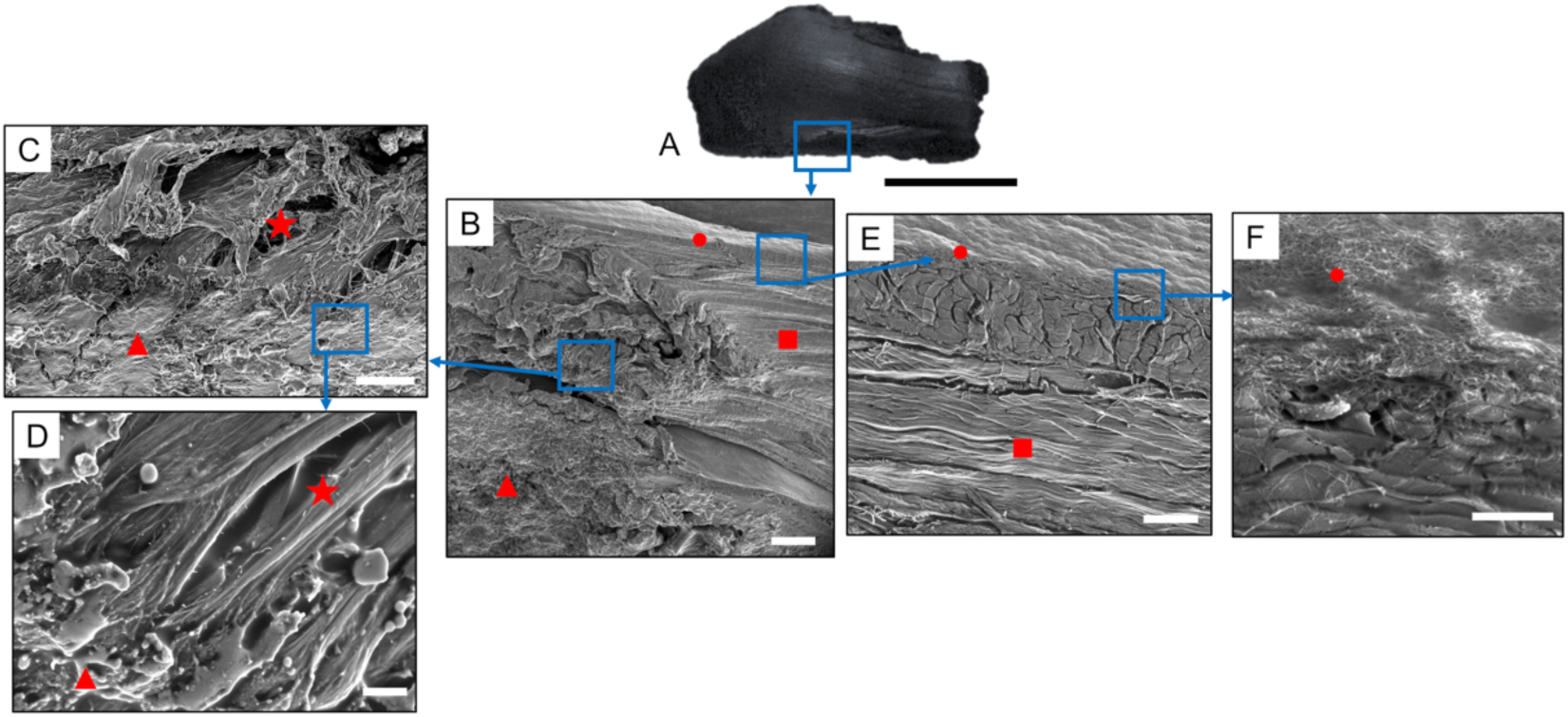
Morphology of cross-section of enthesis over the calcaneal bone of TTSM. SEM images of the TTSM enthesis cross-section: A) picture of the half section of TTSM enthesis (scale bar = 8 mm); B) section of TTSM enthesis with the bone tissue (red triangle), tendon fascicles (red square) and top random fibrils layer (red circle) (magnification 23x, scale bar =500 μm); C) enthesis fascicles with their typical conical junction (red star) entering inside the bone tissue (red triangle) (magnification 5000x, scale bar =5 μm); D) zoom-in at the conical junction of enthesis (red stars) where fibrils of the mFC enter into the bone (red triangle) (magnification 1500x, scale bar =10 μm); E) image tendon fibrils and fascicles (red squares) and top random fibrils layer (red circle) (magnification 90x, scale bar =50 μm); F) zoom-in at the interface between tendon tissue (red square) and random fibrils layer (red circle) (magnification 5000x, scale bar =5 μm).

### 3.2. Histological investigation

The safranin-O staining of the sheep TTSM tendon-to-bone unit in combination with the fast green allowed for the visualization of the histological features of the enthesis (Fig. 5). Safranin-O stains glycosaminoglycans (GAGs) and proteoglycan–rich areas of bright orange or red color while fast green stains collagen-rich areas of green color. The staining allowed to clearly visualize the tendon, enthesis and bone sides for the TTSM tendon-to-bone interface. Both the tendon and bone sides were highlighted in green while the fibrocartilaginous transition between the two tissues appeared stained in shades of purple, red and orange due to the high content of GAGs and proteoglycans at the enthesis region (Fig. 5A). Furthermore, this staining allowed us to highlight the fibrous structure of the epitenon, the protective fibrous sheath made of randomly arranged collagen fibrils protecting the tendon tissue. In particular, the epitenon protects the tendon from abrasive damage while keeping the tissue lubricated during movements. In sheep TTSM tendon-to-bone samples, the epitenon was observed to become visibly thicker, consistently with the SEM images of Fig. 4, at the enthesis and the first centimeters of the bone side (Fig. 5B) before it transitioned to a more cartilage-like tissue over the bone, creating a low-friction lane to accommodate the SDFT (Fig. 5C).

**Fig. 5.**
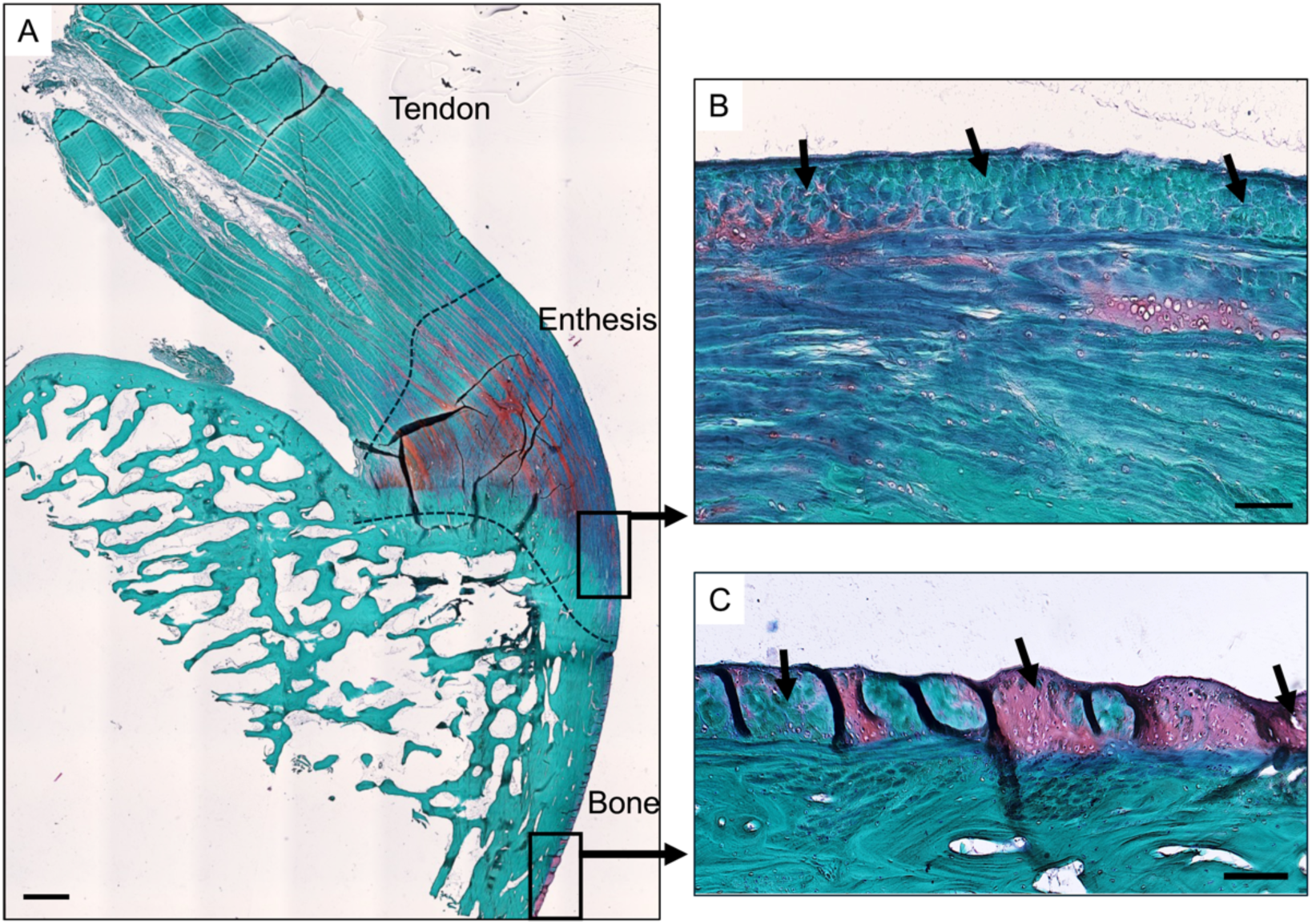
Safranin-O staining of the sheep Achilles tendon-to-bone unit. A) Overview of the tendon-to-bone unit. Dashed lines indicate the respective transitions from tendon to enthesis to bone (scale bar = 1000 µm). B) Zoomed-in view of the epitenon. Black arrows highlight the fibrous morphology of the thick tendon sheet observable along the surface of the enthesis (scale bar = 100 µm). C) Zoomed-in view of the surface of the bone tissue. Black arrows highlight the thick layer of fibrocartilage coating the surface of this region (scale bar = 100 µm).

### 3.3. qBEI and Nanoindentation

A qualitative analysis of the qBEI revealed that the mFC region was more mineralized than the adjacent subchondral bone, with several regions having calcium content even exceeding 27 wt% (Fig. 6). The nanoindentation at the enthesis side highlighted that mFC was also stiffer and harder than bone (in agreement with a higher mineral content). At the interface between non-mineralized and mineralized tissues, steep gradients in mineralization, indentation modulus and hardness were observed. Specifically, the mineral content increased from 5 to 27 wt%, the indentation modulus from 5 to 30 GPa and the hardness from 0.2 to 1.2 GPa when crossing the n-mFC/mFC interface over a thin region of about 10-15 µm. Both mineral content and mechanical properties showed a small decrease when entering in bone. In Fig. 6D, former tidemarks indicating the position of the mineralization front were visible and directly reflected in spatial variations of mechanical properties (Fig. 6G-J). Concerning the trabecular bone (Fig. 6K-6N), the results observed in the indentation modulus evaluation (Fig. 6M) were in the range of 15-22 GPa and the indentation hardness (Fig. 6N) in the range of 0.5-0.8 GPa.

**Fig. 6.**
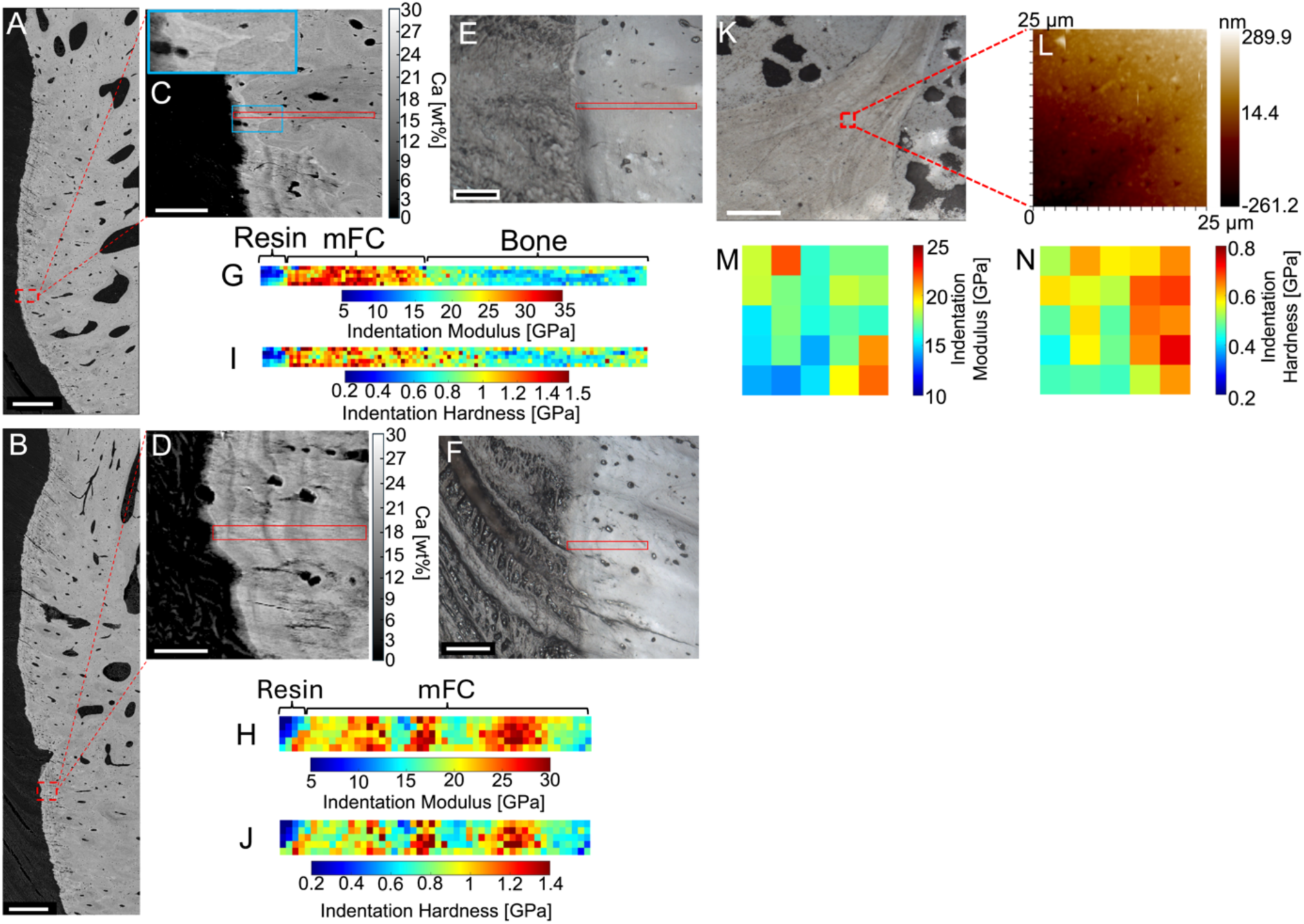
qBEI and nanoindentation of sheep calcaneal bone, TTSM enthesis and trabecular bone: A, B) qBEI overviews of the enthesis region of two different specimens (scale bar = 500 μm); C, D) close up on the qBEI maps of the indented area (red boxes), gray level associated with calcium content (Ca [wt%]) (C: scale bar = 100 μm, D: scale bar = 50 μm); E, F) corresponding optical micrographs showing the indented areas (red boxes) and confirming the enthesis location (tendon = left side, bone = right side) (scale bar = 90 μm); G, H) 2D colormaps of indentation modulus; I, J) 2D colormaps of indentation hardness; K) nanoindenter optical microscope view of the indented region (in red) in trabecular bone (scale bar = 90 μm); L) Scanning probe microscope of the indented region showing the roughness; M) colormap of indentation modulus in trabecular bone; N) colormap of indentation hardness in trabecular bone.

### 3.4 Mechanical tensile tests and DIC

To elucidate the mechanical tensile performances of TTMS and how the collagen fibrils ECM heterogeneity (see Fig. 3-5) influences the superficial strain distribution during the application of the load, mechanical tensile tests coupled with DIC were carried out. The mean ± SD force and stress on the percentage of failure strain curves of the different specimens are shown in (Fig. 7A and 7B) and summarized in Table 1. A representative force-strain curve of a specimen is shown in Fig. 7D.

**Fig. 7.**
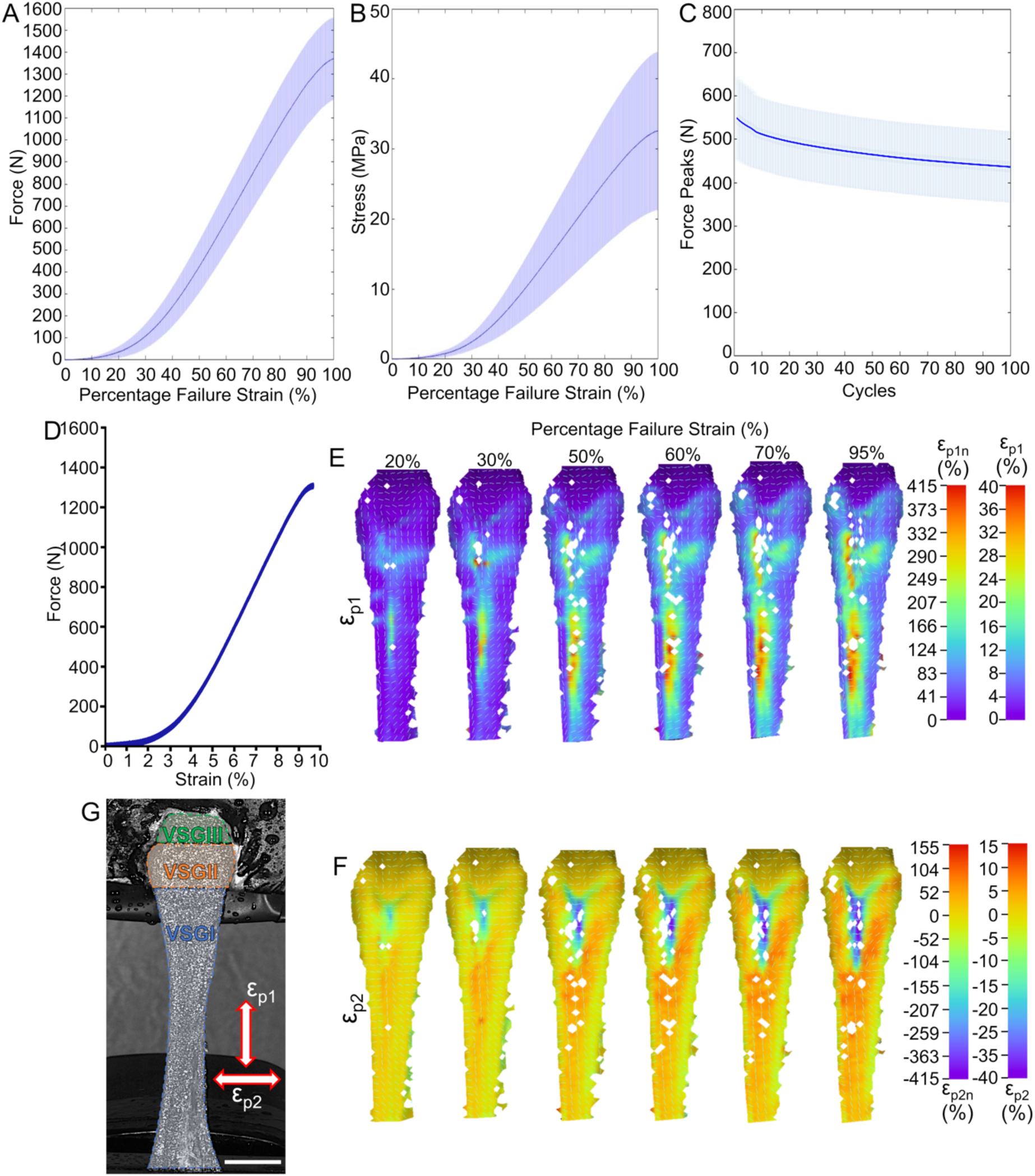
Mechanical characterization and superficial strain distribution of TTSM. A) Mean force on the percentage of failure strain curve of samples (solid line = mean; light lines = SD); B) mean stress on percentage of failure strain curve of samples (solid line = mean; light lines = SD); C) mean peaks of force of samples during the cyclic tests (solid line = mean; light lines = SD). D) Typical force-strain curve of the specimen during the DIC tensile test where: E) represents the 3D view of ε_p1_ at different percentages of failure strain (color bars show the values of strain concentration factor ε_p1n_ and ε_p1_) while F) represent the 3D view of ε_p2_ at the same percentages of failure strain (color bars show the values of strain concentration factor ε_p2n_ and ε_p2_). G) Representation of the virtual strain gauges drawn on a typical specimen (scale bar = 10 mm).

All samples showed the typical non-linear behavior with a toe region extended up to 3 - 3.5 % of nominal strain. Afterward, the curves exhibited a linear behavior, reaching values of failure force of F_F_ = 1369 ± 187 N (σ_F_ = 33±11 MPa) at a failure strain of ε_F_ = 8.2 ± 1.0 %. The stiffness of TTSM samples was K = 309 ± 38 N/mm, resulting in mean values of elastic modulus of E = 531 ± 112 MPa. These values allowed the tendons to store mean values of work to failure of W_F_ = 0.0012 ± 0.0006 J/mm^3^ and a related energy of J_F_ = 3520 ± 850 N/mm. The inflection point strain resulted higher compared with the physiological 5 % of strain used for the later cyclic tests IFε = 7.8 ± 1.8 % (IF_%εF_ = 86 ± 11 %). Parallelly, the inflection point force and strain were IFF = 84 ± 18 N and IFσ = 28 ± 14 MPa (mechanical properties of the single samples are reported in Table S1).

DIC technique provided a deep insight into understanding the local strain concentration effects occurring in the different regions of samples (Fig. 7, Table S2 - S4, Video S1, S2). The systematic and random errors were 0.18% and 0.14%, respectively (Table S2). A representative DIC analysis of a typical specimen is shown in (Fig. 7E-7F) where the evolution of ε_p1_ (ε_p1n_) and ε_p2_ (ε_p2n_) is illustrated at distinct values of the percentage of failure strain. Over the bone (RVI, Fig. 3A), both ε_p1_ and ε_p2_ were approximately close to 0%, indicating that this anatomical district did not substantially contribute to the elastic mechanical behavior of tested samples. In contrast, the tendon side (RI, Fig. 3A) exhibited larger incremental strain during the tensile tests (Fig. 7E), approaching local maximum values of ε_p1_ in the range of 32 - 40% along the centerline of the tendon tissue.

In addition, RII, RIV, and RV exhibited (i.e. enthesis region) an ε_p1_ behavior intermediate between the two opposite regions over the bone (RVI) and the tendon (RI) tissue, accomplishing a progressive transition of deformation that reached values of 10 -18% (ε_p1n_ = 103 - 186%) at 95 % of failure strain.

Interestingly ε_p2_ was consistently positive along RI, the sides of RII, RIV and RV with values ranging from 5% up to 10% (ε_p2n_ = 52 - 103 %) highlighting the auxetic macroscopic behavior of the tendon and enthesis tissue. The region in between the enthesis (i.e. the center of RII and RIII) exhibited a progressive increase of ε_p1_, reaching ranges from 8 – 10 % (ε_p1n_ = 82 - 103 %) of strain. As demonstrated by the high negative level of ε_p2_ approached in the center of RII and in RIII (from -10% up to -40% at 95% of failure strain) these regions contracted significatively during loading.

Considering these findings, the majority of strain seems to be mostly concentrated in the tendon and enthesis region (region I-V), highlighting their distinct behavior compared with RVI. The need to furnish a strain distribution overview along the whole TTSM (including the enthesis and bone regions) while preserving the sample inside the field of view of the DIC system during the tensile test, allowed to clearly highlight the transitions zones between RI-RVI but reduced the accuracy of the strains measurements.

To allow the calculation of averaged ε_p1_ and ε_p2_ values, three virtual strain gauges were used. As visible from Table S3 and S4, the mean strain values of all samples at the different percentages of apparent strain, as well as their maximum and minimum peaks, were mostly in accordance, for the bone (VSGIII) and tendon (VSGI) region (i.e. at 95% of failure strain: VSGIII: ε_p1_mean_ = 0.41 ± 0.36 %, ε_p1n_mean_ = 4.29 ± 3.10 %, ε_p2_mean_ = -0.12 ± 0.08 %, ε_p2n_mean_ = -1.39 ± 0.98; VSGI: ε_p1_mean_ = 7.68 ± 3.46 %, ε_p1n_mean_ = 84.59 ± 38.75 %; ε_p2_mean_ = 4.07 ± 2.15 %, ε_p2n_mean_ = 43.90 ± 20.97 %), with the typical strain ranges of the specimen described above (Fig. 7E, 7F). Conversely for the enthesis, having included in the virtual strain gauge all the regions RII-RV, their mean values masked the auxetic behavior the sides of RII and RIV-RV regions due to presence of extend compressive zone in the center of RII and RIII (i.e. at 95% of failure strain: VSGII: ε_p1_mean_ = 7.08 ± 2.41 %, ε_p1n_mean_ = 78.56 ± 27.15 %, ε_p2_mean_ = -1.17 ± 2.77 %, ε_p2n_mean_ = -13.95 ± 29.82 %).

### 3.5 Cyclic tests

The cyclic tests carried out on TTSM (Fig. 7C) revealed a progressive loss of force from the first (F_P_ = 549 ± 95 N) up to the hundredth cycle (F_P_ = 436 ± 82 N) with a mean force loss in percentage at the last package of 21 ± 7 %. The higher mean force loss between consecutive packages (10 cycles each) happened in the first four (F_packages_loss_P1vsP2_ = 3.2 ± 1.0 %; F_packages_loss_P2vsP3_ = 2.2 ± 0.7 %_;_ F_packages_loss_P2vsP3_ = 1.8 ± 0.6 %; F_packages_loss_P3vsP4_ = 1.5 ± 0.6 %) reaching a progressive stabilization up to the last three packages (F_packages_loss_P8vsP9_ = 0.9 ± 0.4 %; F_packages_loss_P9vsP10_ = 0.8 ± 0.3 %) (Table 2 and Table S5).

**Table 2.** Mean ± SD of: the force peaks (F_P_) values, the percentage of the force loss between the first cycle’s peaks of the first packages compared with the first cycle’s peak of each packages (F_Peak_loss_) and the percentage of force peak loss (F_Packages_loss_) between the last cycle’s peaks of two consecutive packages (10 cycles) of the cyclic tests of TTSM samples.

| Samples | Package |  |  |  |  |  |  |  |  |  |
| --- | --- | --- | --- | --- | --- | --- | --- | --- | --- | --- |
| (n = 6)<br>Cycle | P1 | P2 | P3 | P4 | P5 | P6 | P7 | P8 | P9 | P10 |
| $F_P$<br>(N) | 549±95 | 512±84 | 495±84 | 485±84 | 473±83 | 465±83 | 457±83 | 451±83 | 446±83 | 441±83 |
| $F_{\text{Peak\_loss}}$<br>(%) | 7±2 | 10±3 | 12±4 | 14±4 | 15±5 | 17±5 | 18±6 | 19±6 | 20±7 | 21±7 |
|  | P1 vs P2 | P2 vs P3 | P3 vs P4 | P4 vs P5 | P5 vs P6 | P6 vs P7 | P7 vs P8 | P8 vs P9 | P9 vs P10 | - |
| $F_{\text{Packages\_loss}}$<br>(%) | 3.2±1.0 | 2.2±0.7 | 1.8±0.6 | 1.5±0.6 | 1.4±0.6 | 1.2±0.5 | 1.0±0.4 | 0.9±0.4 | 0.8±0.3 | - |

## 4. Discussion

In this study, the complementary use of different imaging techniques, such as SEM and histology, coupled with mechanical characterizations monitored with DIC, allowed a comprehensive description of the structure and mechanics of the sheep TTSM-calcaneal bone complex. This tendon, along with the SDFT one, constitutes the ovine Achilles tendon. Due to the mechanical relevance and the similarity of the ECM composition between sheep and human tendons, [52,53], this animal model is often adopted in preclinical *in vivo* studies to test promising scaffolds and implantable devices before moving to clinical trials. In contrast to the SDFT that slides over the TTSM reaching the bone at the foot [38], the TTSM connects directly with the calcaneal bone at the Achillis level [38], producing a well-defined enthesis, making it an excellent site for investigating the mechanics of devices meant to treat human enthesis. Looking at the nanostructure of TTSM ECM, SEM investigation revealed that collagen fibrils had a close morphological similarity in terms of organization, orientation, and thickness with the human counterpart [7,8,54] (Fig. 1A and Fig. 3, 4). Although macroscopically tendon tissue seems to attach to the calcaneal bone preserving its morphology and structure (Fig. 1B), SEM images of TTSM enthesis revealed that fibrils progressively halved their cross-section increasing their randomicity in a region-specific manner (Fig. 3). In the internal cross-section of the same area (Fig. 4) instead, TTSM fascicles confirmed the typical conical shape of enthesis at the n-mFC level (Fig. 4C - 4D) [55]. The superficial fibrillar layer above them (Fig. 3A-3I, 4), probably is a local thickening of the epitenon sheath further confirmed by the histology safranin-O staining (Fig. 5B). The physiological function of this local epitenon thickening is likely to provide protection and lubrication to TTSM reducing friction and wear caused by the continuous sliding on it of the SDFT [38].

The results of nanoindentation of the mFC were fully in accordance with the previous literature concerning large animals and humans [56]. Here indeed, the high mechanical stress/strain that stimulate this anatomical site produces an up-regulation in the production of mineralization. This over mineralization results in a narrow transition region (10-15 μm in width) with a gradient of mineralization (Fig. 6A-6J). Conversely, in small animals such as rats, the amount of mineralization is lower due to the lower forces involved and the different anatomical conformation [11,57]. Similarly, the indentation modulus and hardness found on the trabecular bone reached average values observed in the literature [58] (Fig. 6K-6N).

Mechanically speaking, TTSM showed the typical nonlinear behavior of T/L tissues reported in literature [54]. The DIC investigation during tensile tests allowed to clarify the superficial strain distribution of such complex anatomical district. Along with the progression of tensile tests, ε_p1_ and ε_p2_ allowed to highlight the typical superficial strain distribution of tendons and ligaments reported in literature [20–23], especially when considered as mean values with the virtual strain gauges. The concentration of high peaks of ε_p1_ (ε_p1n_ = 331 - 414%) along the tendon (RI), suggested that this region is the main responsible of the elasticity of samples. Indeed, these values were consistently higher than the 9.2% of nominal strain (i.e. 95% of failure strain Fig. 7E). In RI the values of ε_p2_ were the highest approaching local peaks up to 15% (ε_p2n_ = 155%) highlighting the auxetic behavior of sheep TTSM tissue in accordance with reports in literature on human and animal T/L [20–23]. Moreover, the particular “Y” shape distribution of ε_p2_ suggested that TTSM is internally composed by two major tertiary fiber bundles. This anatomical strategy is probably functional to macroscopically magnify the stress concertation reducing effect that the tendon/enthesis fascicles of TTSM do at the microscale, when inserting in the bone tissue with their typical conical shape [15]. In fact, the auxetic expansion of the enthesis tissue both macroscopically [23] and microscopically [15] detected in literature is functional to locally enlarge the cross-section of the T/L under load, to consequently reduce the values of concentrated stress in that area. Moreover, the auxetic expansion of the tendon (RI), the body (RIV) and the sides (RV) of enthesis, produced a consecutive compression in the regions below (RII) and in the center (RIII) of the enthesis, probably functional to compensate the effect of the surrounding regions.

Tensile tests also confirmed that the inflection point of TTSM resulted in higher strains (i.e. IFε = 7.8 ± 1.8 %, IF_%εF_ = 86 ± 11 %) compared with the typical 5 % of strain used for the later cyclic tests, and often applied in literature to test the performances of scaffolds [50,51,59], validating the suitability of the use up to these strain levels to test the cyclic behavior of biomimetic scaffolds for T/L tissue in large animal models. In fact, after the conclusion of the internal reorganization of the collagen fibrils after the first tens of cycles, TTSM starts losing around or less then 1% of force per each package of 10 cycles. This confirms that working below the inflection point it is possible to preserve the integrity of the majority of collagen fibrils, avoiding the trigger of significant tissue lesion while being below the macroscopic yielding/rupture point. Our work allowed to highlight and confirm several similarities between the TTSM tendon-enthesis-bone chain and human sites including: the morphological similarities of ECM [19], the superficial strain distribution [23] and the mechanics of the small and reduced gradient of mineralization at the enthesis [56].

All the morphological and mechanical findings found in this work will furnish useful suggestions to biofabrication experts working on the design of biomimetic load-bearing scaffolds for the regeneration of the tendon-enthesis-bone chain. At first, to guarantee high mechanical performance of scaffolds (in terms of stiffness, stress and strain values), for both the tendon and bone side, resorbable high-molecular weight bioresorbable polymers, sometimes blended with collagen [50,60–62], have to be considered. To further increase the mechanical strength while achieving a morphological hierarchical biomimicry of the tendon and enthesis sides, the use of biofabrication strategies such as the electrospinning [26,27,63–65] technique and the hierarchical scaffolds [51,66–68], can be considered promising solutions. To mimic the natural gradients of mineralization typical of the enthesis and bone side of the chain, different approaches have been proposed with promising results including gradients of synthetic and natural mineral crystals [26,65]. However, to reproduce such a tiny and abrupt passage from the mFC and the bone tissue as in the TTSM, the merge of multiple biofabrication strategies such as electrospinning and additive manufacturing have offered encouraging preliminary results [69–71].

The study has also some limitations. Only right posterior legs were investigated in this study, making a statistical analysis on the contralateral legs to evaluate possible side-specific differences not possible. Moreover, in the future a higher number of samples for qBEI and nanoindentation shall be investigated to increase the sample size of these investigations. DIC was able to detect the superficial strain distribution of TTSM and the fibrocartilaginous region over the enthesis partially masked the strain evolution of the enthesis fascicles under it. Further investigations by using *in situ* mechanical stepwise tests and DVC shall be carried out to clarify the full-field strain distribution of the enthesis fascicles during the application of a load.

## 5. Conclusions

We carried out a comprehensive morphological, mechanical, and biological characterization aiming at deeply understanding the structure and mechanics of a relevant large animal model such as the sheep TTSM. The SEM morphological characterization showed a superficial abrupt change of fibrillar orientation and dimension by passing from the tendon to the enthesis region. In the internal section of the enthesis region instead, the tendon fascicles attach to the bone tissue with the typical conical shape of non-mineralized fibrocartilage. The histological investigation confirmed that the superficial region of the enthesis was covered by a thick layer of functional cartilage to reduce friction and protect this delicate anatomical site. The mechanical tests coupled with DIC highlighted a progressive decrease of maximum principal strains passing from the tendon to the bone tissue. Moreover, the minimum principal strain confirmed the auxetic mechanical behavior of the tendon tissue with a compressive region at the enthesis caused by the “Y” shape attachment of tendon fascicles to the bone. The qBEI and nanoindentation analyses confirmed respectively higher mineralization and mechanical properties (stiffness and hardness) of the mFC region compared with the cortical and trabecular bone. This study contributed to describe and clarify the relationship between micro-/nanostructure, superficial strain distribution, soft-hard tissue mechanics and step gradient of mineral content/nanomechanics of the tendon-enthesis-bone chain. All these findings will be extremely useful for the biofabrication scientific community interested in designing a new generation of biomimetic hierarchical scaffolds for the regeneration of the tendon and enthesis tissue.

## 6. Acknowledgments

Horizon Europe Marie Skłodowska Curie Fellowship 3NTHESES (n.101061826) is greatly acknowledged for funding the work. I.L.C.O. srl is acknowledged for furnishing the sheep samples. Francesco Vai is acknowledged for the useful suggestions in designing the clamping system for the tensile tests. Luca Cristofolini and Nicola Sancisi are acknowledged for the useful suggestions and the use of their facilities and laboratories for the preparation of samples. Carmen Lopez Iglesias, Hans Duimel, and Willine van de Wetering are acknowledged for their useful advice and help for sample decellularization for SEM investigation. Alessandra Di Lorenzo is acknowledged for his help in the elaboration of mechanical data. We thank Petra Keplinger, Sonja Lueger and Phaedra Messmer for assistance in qBEI measurements. Markus Hartmann and Stéphane Blouin gratefully acknowledge financial support from the Austrian Social Health Insurance Fund (OEGK) and the Research funds of the Austrian workers compensation board (AUVA). Albano Malerba and Alexandra Tits have received funding from the FWO and F.R.S.-FNRS under the Excellence of Science (EOS)-programme (EOS No. 40007553). Luca Raimondi acknowledges the support of Ecosystem for Sustainable Transition in Emilia-Romagna Project, funded under the National Recovery and Resilience Plan (NRRP), Mission 04 Component 2 Investment 1.5— NextGenerationEU, call for tender n. 3277 dated 30 December 2021 Award Number: 0001052 dated 23 June 2022 CUP: B33D21019790006.

## 7. CRediT authorship contribution statement

**A. Sensini**: Writing – original draft, Methodology, Investigation, Visualization, Funding acquisition, Formal analysis, Data curation, Project administration, Conceptualization. **L. Raimondi**: Writing – original draft, Methodology, Investigation, Formal analysis, Data curation. **A. Malerba**: Writing – original draft, Investigation, Visualization, Formal analysis, Data curation. **C. Peniche Silva**: Writing – original draft, Investigation, Methodology, Visualization, Formal Analysis, Data curation. **A. Zucchelli**: Writing – review & editing, Methodology, Resources, Data curation. **A. Tits**: Writing – review & editing, Investigation. **D. Ruffoni**: Writing – review & editing, Resources, Methodology. **S. Blouin**: Writing – review & editing, Investigation, Formal analysis, Methodology, Data curation. **M. Hartmann**: Writing – review & editing, Investigation, Formal analysis, Methodology, Data curation, Resources. **M. van Griensven**: Writing – review & editing, Methodology, Resources, Funding acquisition, Project administration, Conceptualization, Supervision. **L. Moroni**: Writing – review & editing, Methodology, Resources, Funding acquisition, Project administration, Conceptualization, Supervision.

## 8. Declaration of competing interest

The authors declare that they have no known competing financial interests or personal relationships that could have appeared to influence the work reported in this paper.

## 9. Data availability

Data will be made available from the authors under request.

## 10. Supplementary Information

Supplementary information of this article can be found online.

## Supplementary Material

**Video S1.**
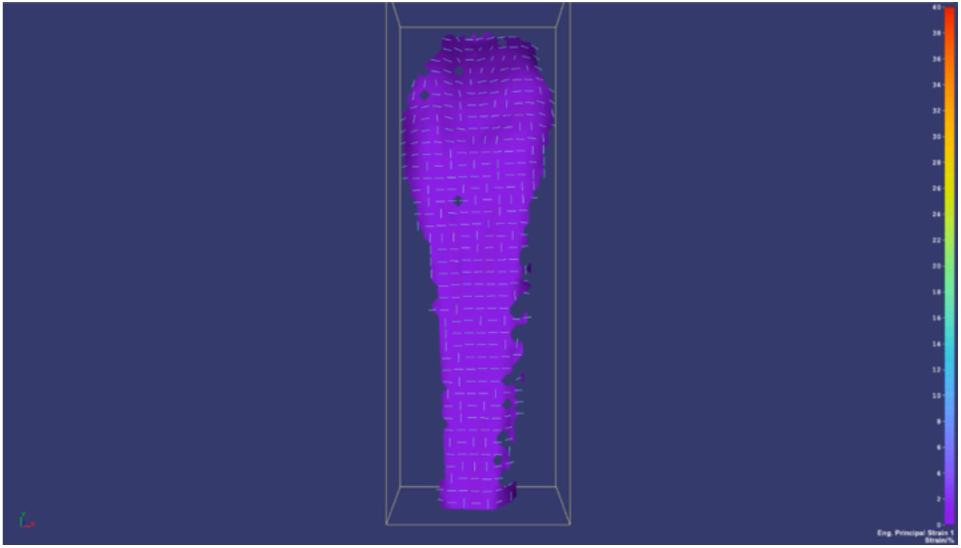
Maximum principal strain (ε_p1_) evolution in the representative sample of Figure.7.

**Video S2.**
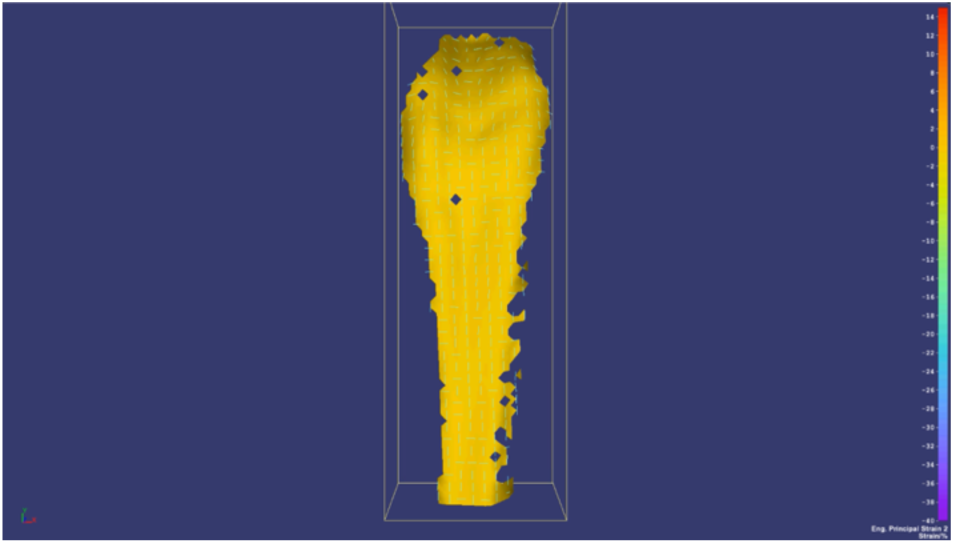
Minimum principal strain (ε_p2_) evolution in the representative sample of Figure.7.

**Table S1.** Mechanical tensile properties of TTSM.

| Sample | $F_F$<br>(N) | $\sigma_F$<br>(MPa) | $\epsilon_F$<br>(%) | E<br>(MPa) | K<br>(N/mm) | $W_F$<br>(J/mm<sup>3</sup>) | IFF<br>(N) | IFσ<br>(MPa) | IFε<br>(%) | IF$\% \epsilon_F$<br>(%) |
| --- | --- | --- | --- | --- | --- | --- | --- | --- | --- | --- |
| ST1 | 1308 | 34 | 9.7 | 546 | 299 | 0.0012 | 1183 | 31 | 8.7 | 90 |
| ST2 | 1221 | 29 | 8.4 | 515 | 294 | 0.0009 | 615 | 14 | 5.5 | 66 |
| ST3 | 1363 | 22 | 8.0 | 398 | 346 | 0.0007 | 1200 | 20 | 6.9 | 86 |
| ST4 | 1671 | 54 | 11.1 | 734 | 323 | 0.0024 | 1657 | 54 | 10.9 | 98 |
| ST5 | 1161 | 27 | 8.4 | 506 | 246 | 0.0008 | 934 | 21 | 7.0 | 83 |
| ST6 | 1491 | 30 | 8.5 | 484 | 348 | 0.0011 | 1460 | 29 | 8.0 | 94 |
| Mean±SD | 1369±187 | 33±11 | 8.2±1.0 | 531±112 | 309±38 | 0.0012±0.0006 | 84±18 | 28±14 | 7.8±1.8 | 86±11 |

**Table S2.** DIC uncertainty (zero-strain) analysis and systematic and random errors.

| Specimen | $\epsilon_{p1\_mean\_zero-strain}$<br>(%) | $\epsilon_{p1\_SD\_zero-strain}$<br>(%) | $\epsilon_{p2\_mean\_zero-strain}$<br>(%) | $\epsilon_{p2\_SD\_zero-strain}$<br>(%) |
| --- | --- | --- | --- | --- |
| ST1 | 0.01 | 0.1 | -0.01 | 0.07 |
| ST2 | 0.04 | 0.18 | 0.00 | 0.20 |
| ST3 | 0.10 | 0.15 | 0.04 | 0.14 |
| ST4 | 0.02 | 0.12 | -0.07 | 0.12 |
| ST5 | 0.07 | 0.09 | 0.35 | 0.08 |
| ST6 | 0.50 | 0.20 | 1.13 | 0.17 |
| Mean (%) | 0.12 | - | 0.24 | - |
| Quadratic mean (%) | - | 0.15 | - | 0.14 |
| Systematic Error (%) | 0.18 | - | - | - |
| Random Error (%) | - | 0.14 | - | - |

**Table S3.**
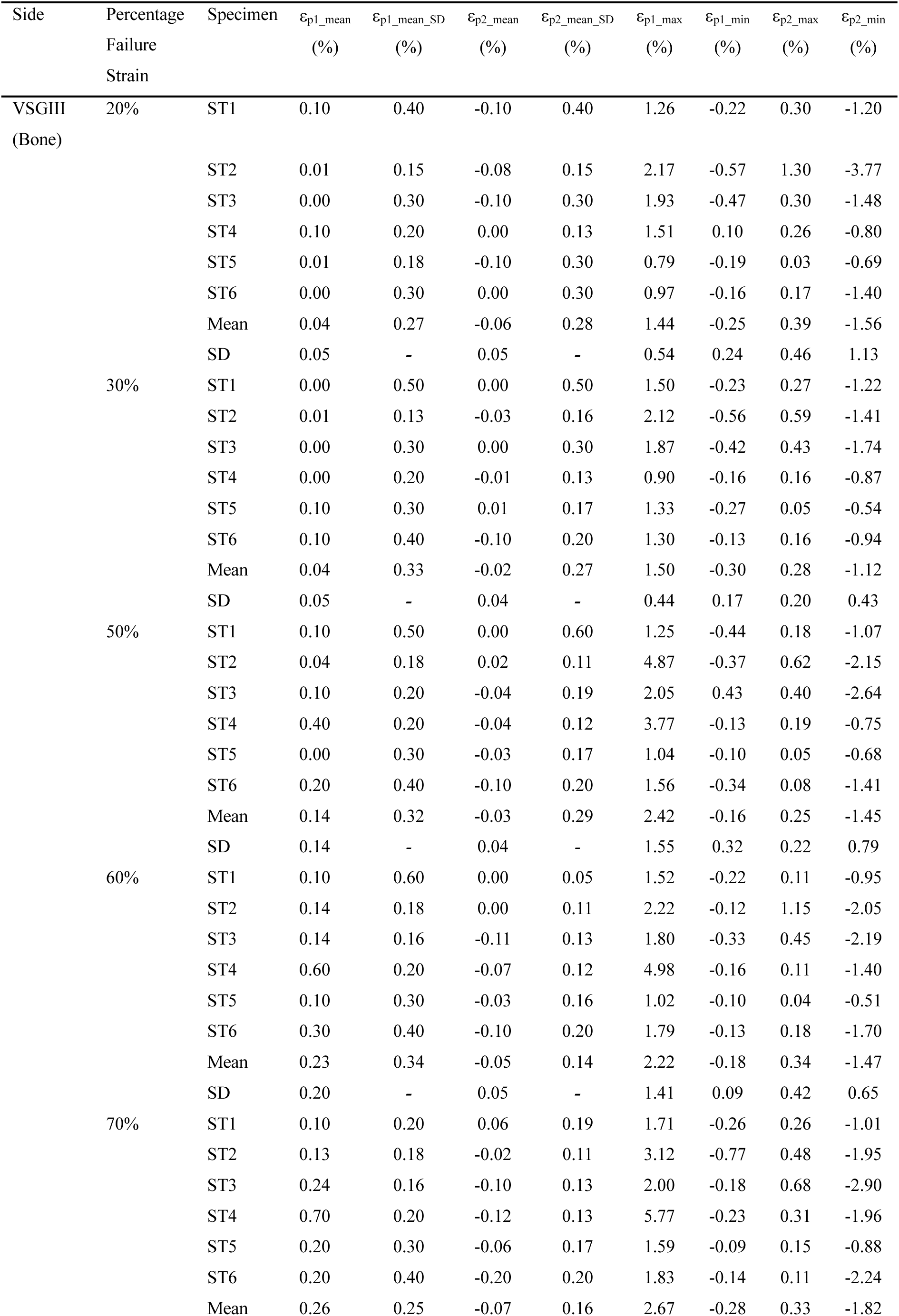
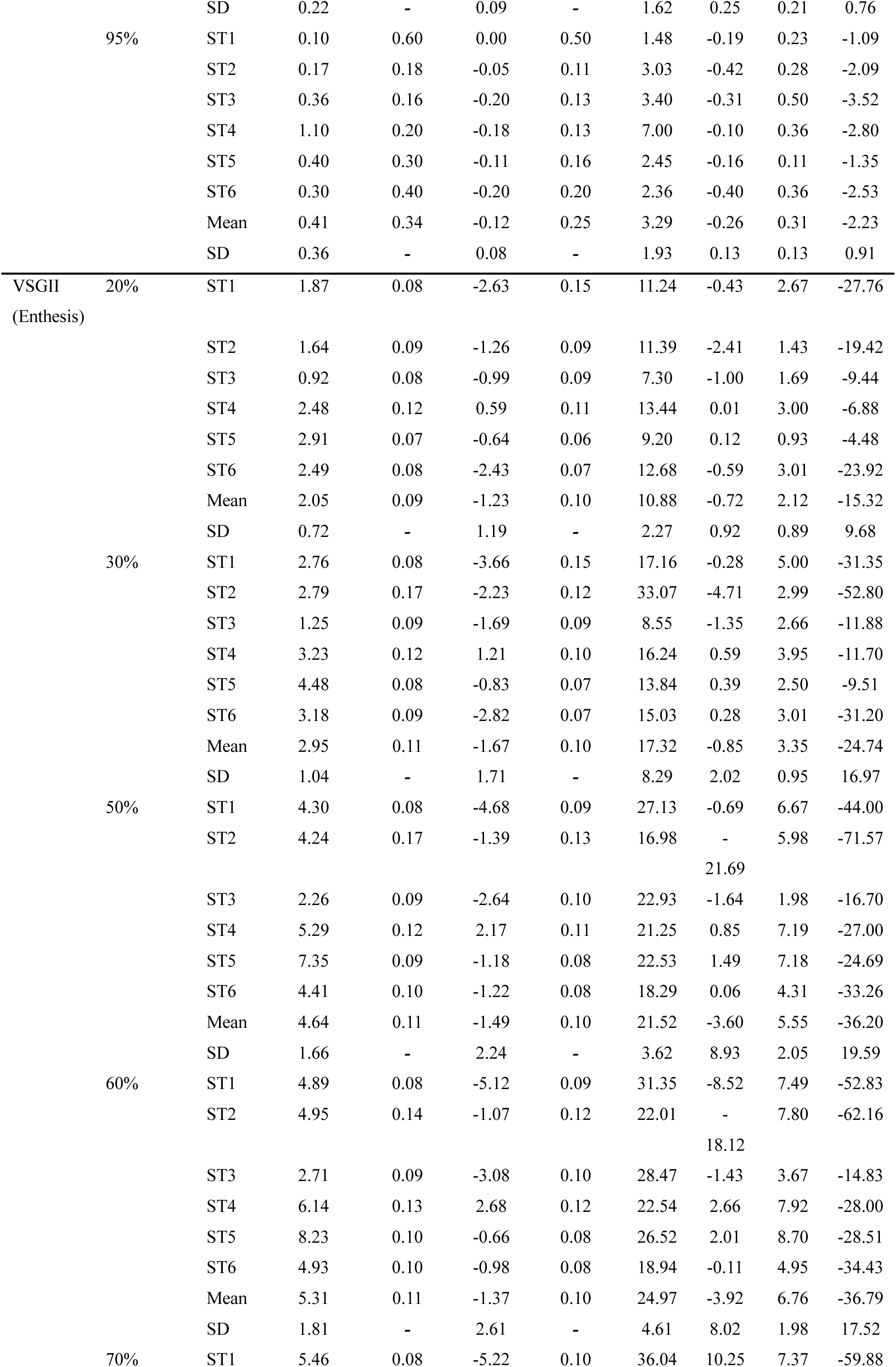
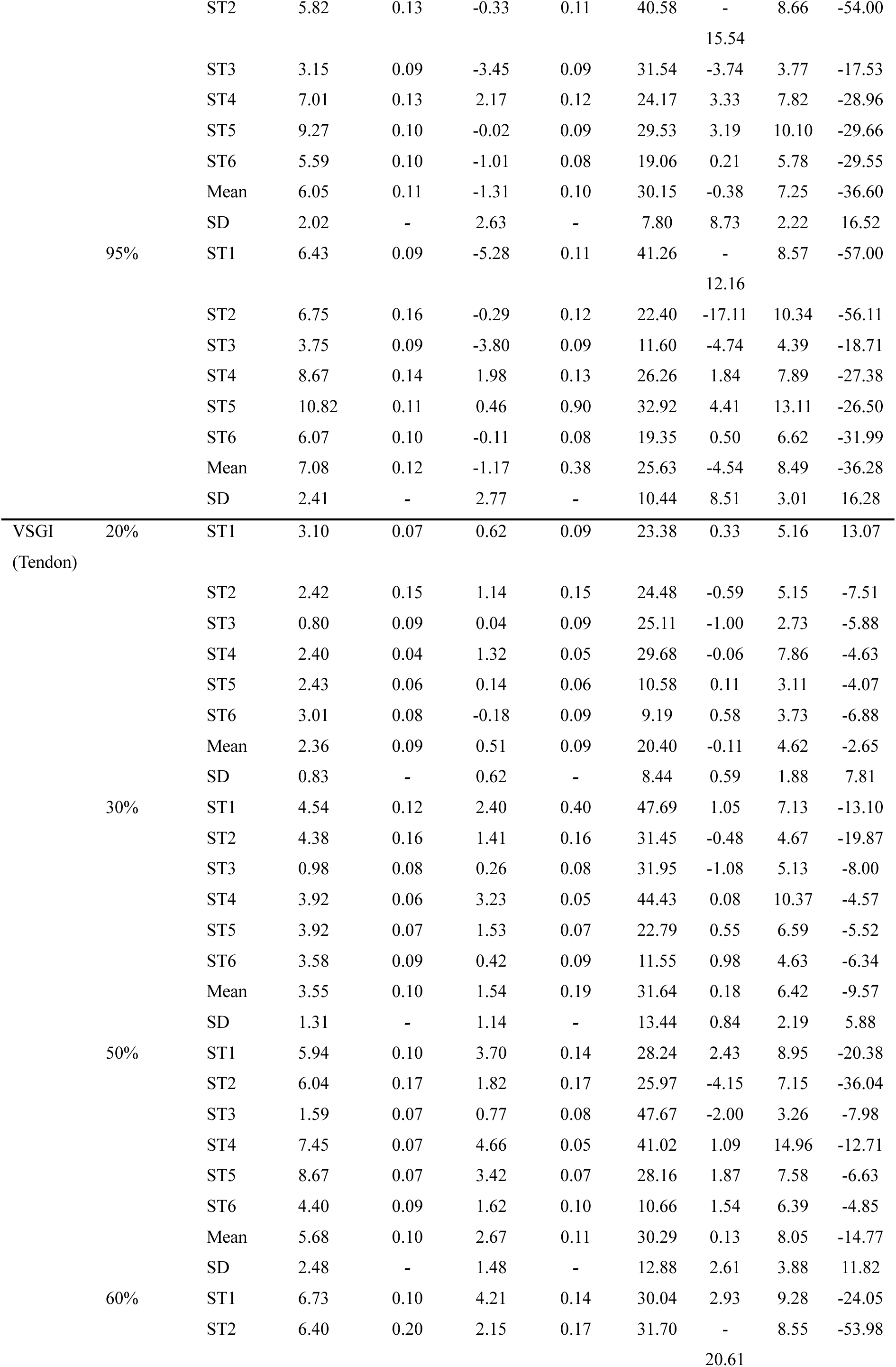
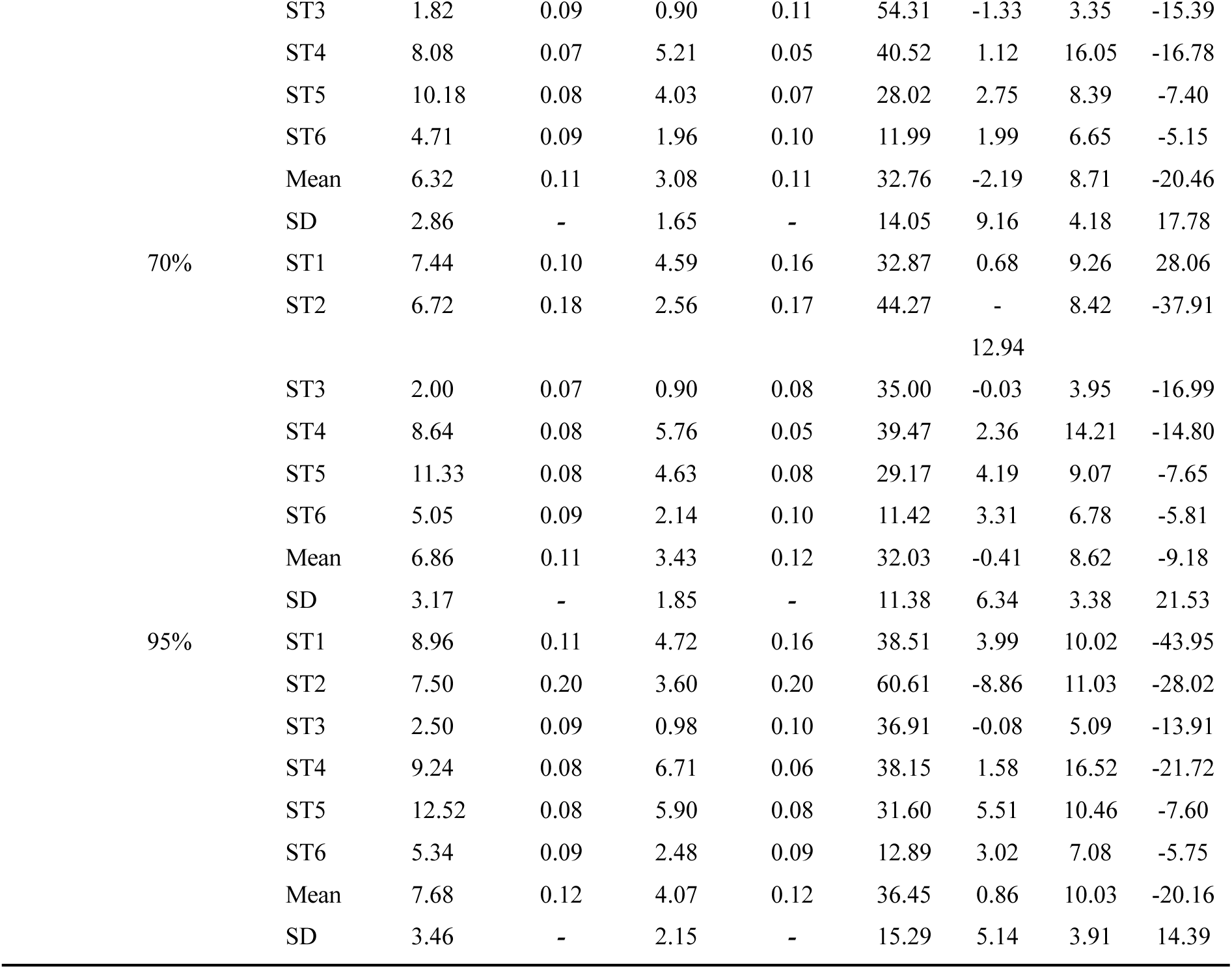
Principal strains (ε_p_) at the different percentage of failure strain steps in the bone, enthesis and tendon sections of TTSM specimens.

**Table S4.**
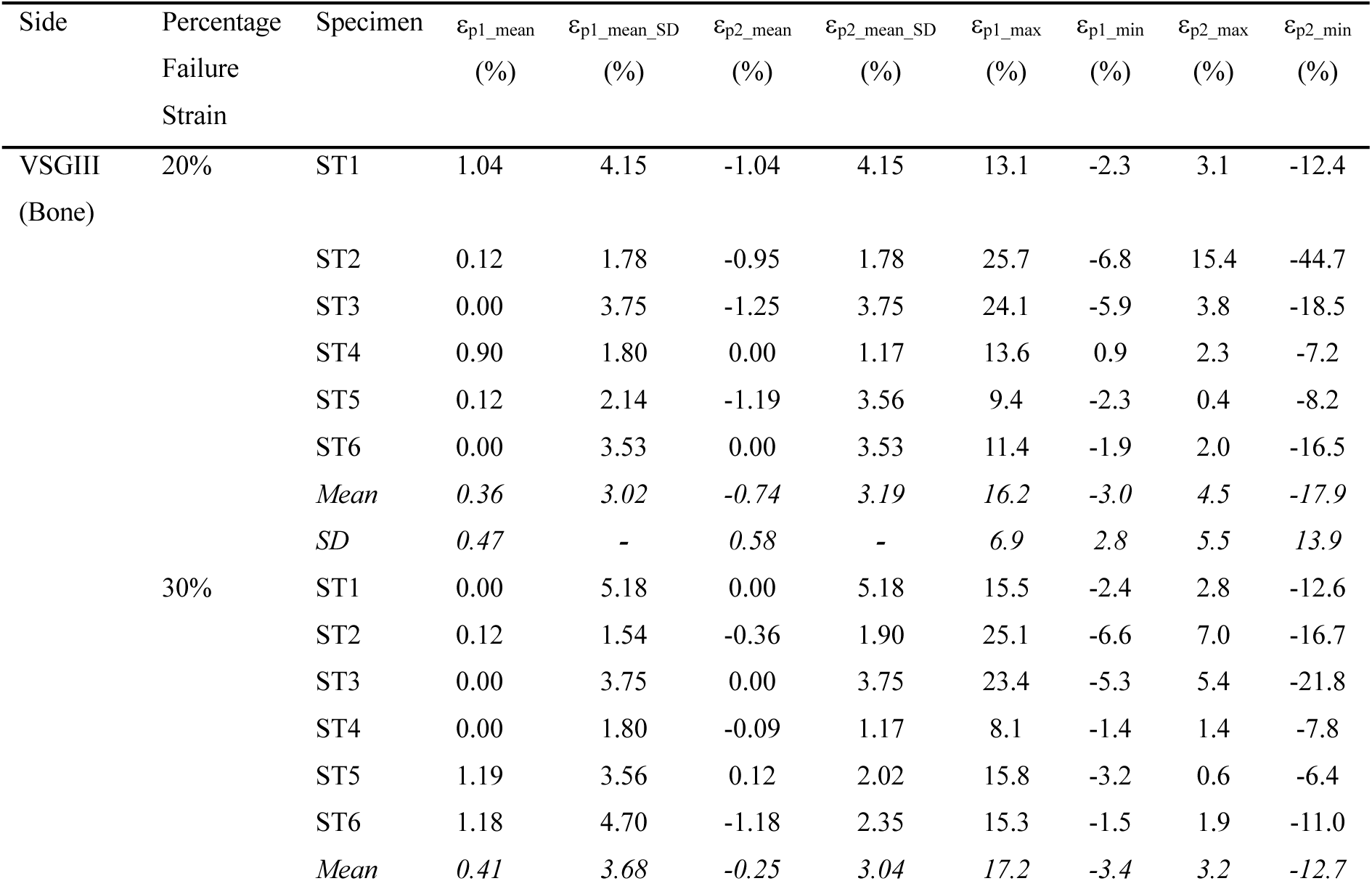
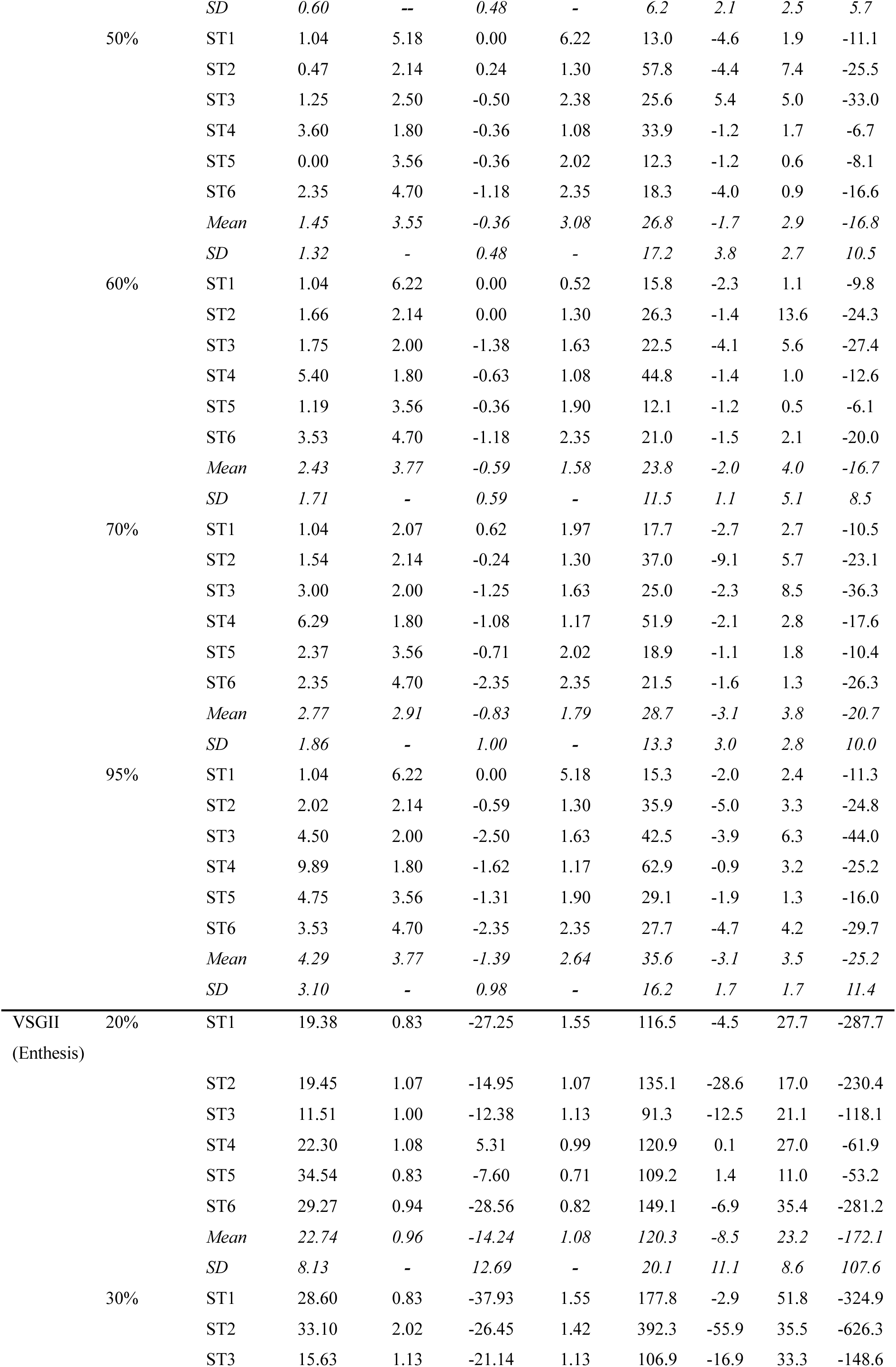
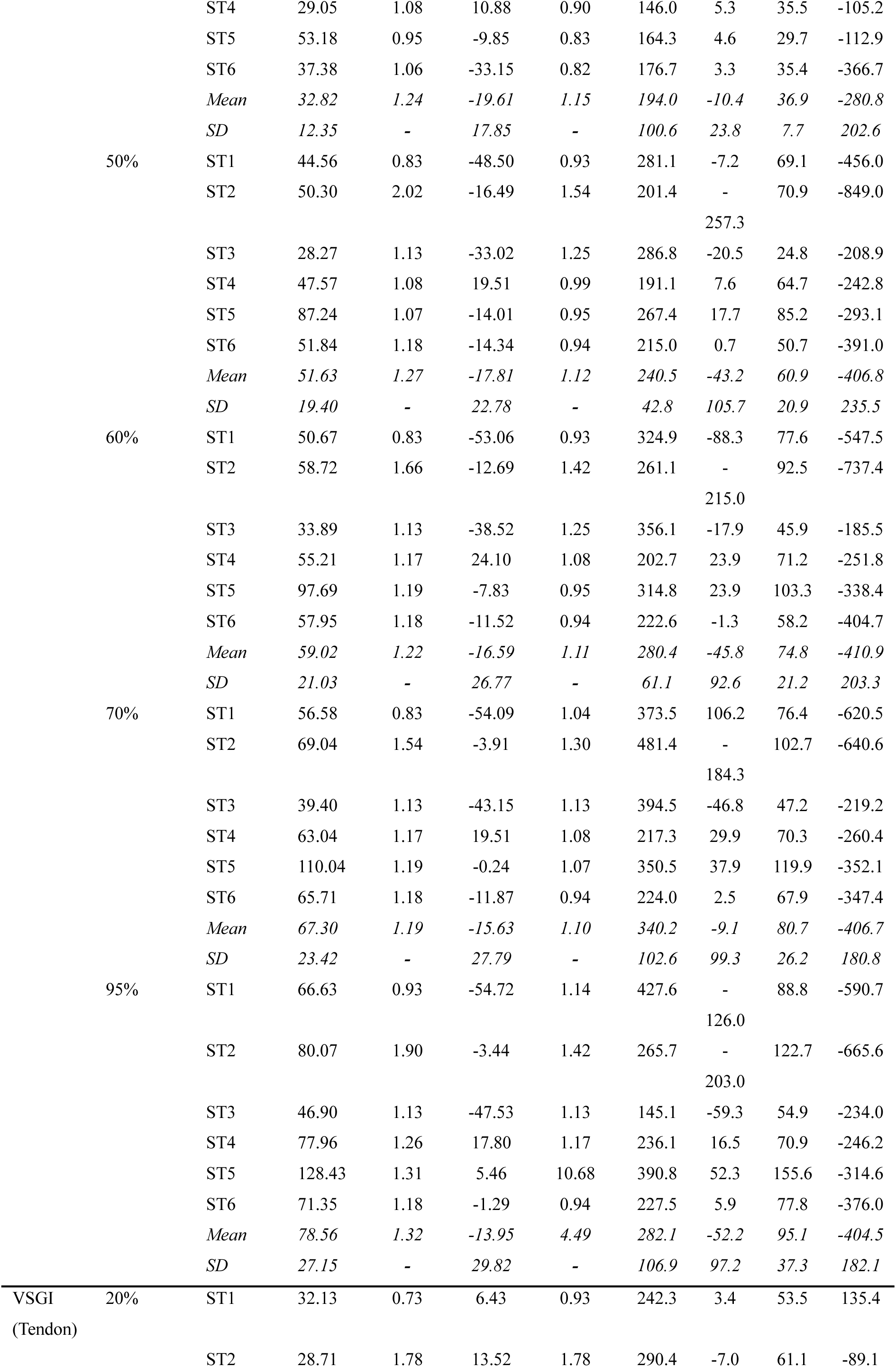
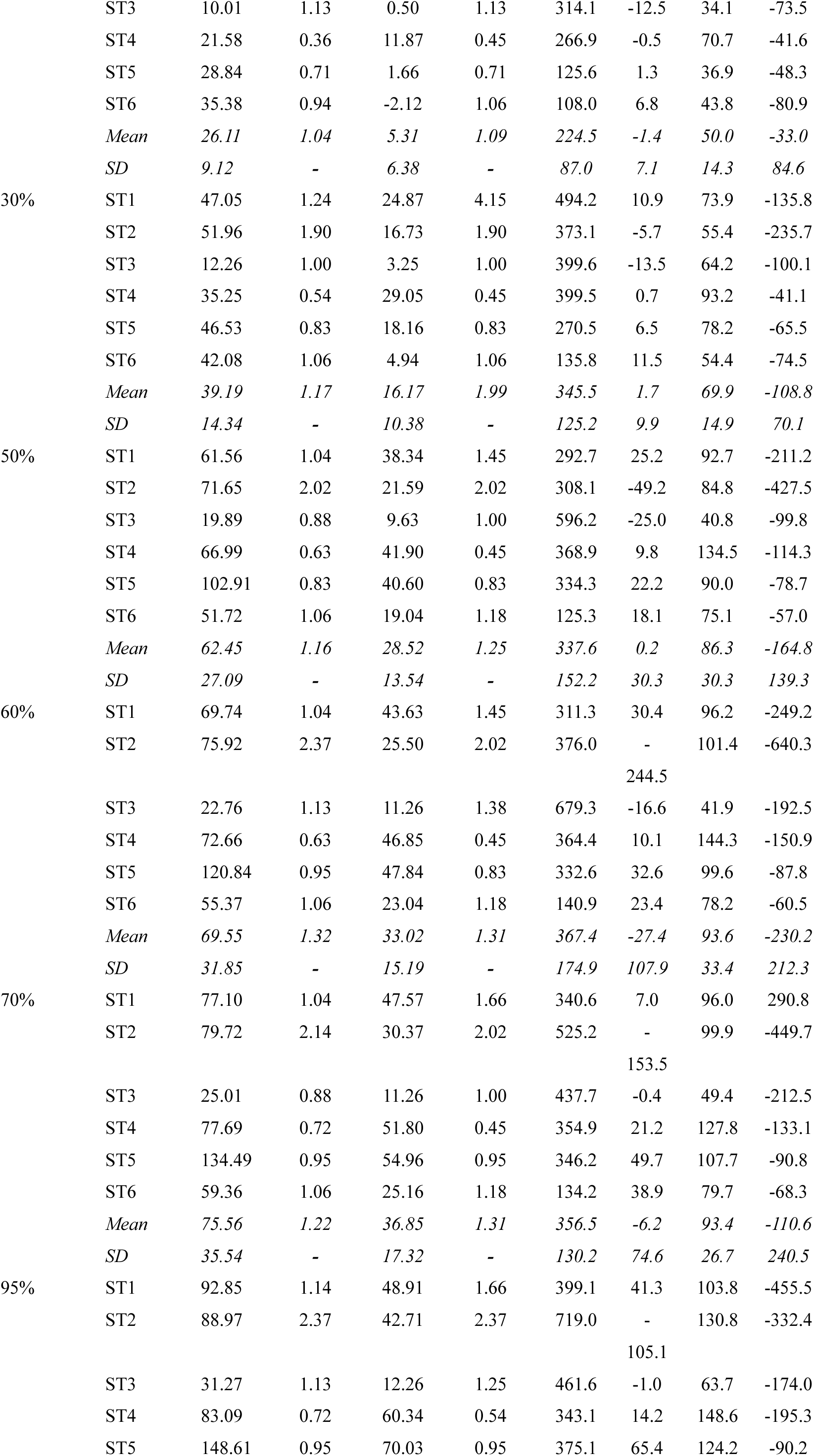
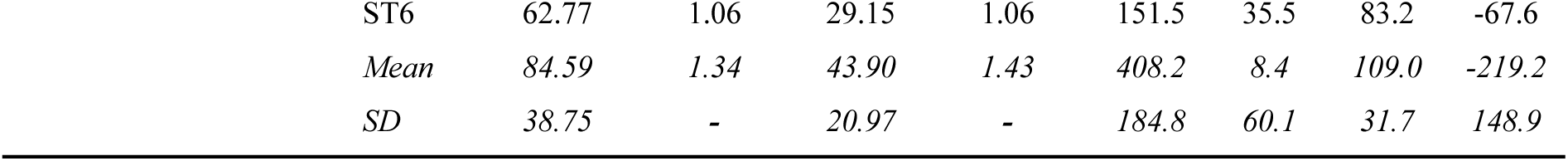
Principal strains normalized (ε_pn_) at the different percentage of failure strain steps in the bone, enthesis and tendon sections of TTSM specimens.

**Table S5.** Values of single samples and mean ± SD of: the force peaks (F_P_) values, the percentage of the force loss between the first cycle’s peaks of the first packages compared with the first cycle’s peak of each packages (F_Peak_loss_) and the percentage of force peak loss (F_Packages_loss_) between the last cycle’s peaks of two consecutive packages (10 cycles) of the cyclic tests of TTSM samples.

|  | Sample | Package |  |  |  |  |  |  |  |  |  |
| --- | --- | --- | --- | --- | --- | --- | --- | --- | --- | --- | --- |
|  | Cycle | P1 | P2 | P3 | P4 | P5 | P6 | P7 | P8 | P9 | P10 |
| $F_P$<br>(N) | SC1 | 390 | 365 | 350 | 341 | 331 | 324 | 317 | 311 | 390 | 365 |
|  | SC2 | 551 | 515 | 490 | 476 | 460 | 450 | 441 | 434 | 551 | 515 |
|  | SC3 | 556 | 505 | 483 | 471 | 456 | 445 | 434 | 425 | 556 | 505 |
|  | SC4 | 570 | 550 | 541 | 536 | 529 | 526 | 522 | 520 | 570 | 550 |
|  | SC5 | 689 | 622 | 603 | 592 | 575 | 563 | 552 | 543 | 689 | 622 |
|  | SC6 | 538 | 514 | 500 | 493 | 486 | 481 | 477 | 473 | 538 | 514 |
| | Mean±<br>SD | 549±95 | 512±84 | 495±84 | 485±84 | 473±83 | 465±83 | 457±83 | 451±83 | 446±83 | 436±82 |
| $F_{Peak\_loss}$<br>(%) | SC1 | 6.46 | 6.64 | 9.16 | 3.41 | 9.72 | 4.56 | 6.46 | 6.64 | 9.16 | 3.41 |
|  | SC2 | 10.2 | 11.1 | 13.1 | 5.05 | 12.4 | 7.12 | 10.2 | 11.1 | 13.1 | 5.05 |
|  | SC3 | 12.8 | 14.2 | 15.8 | 6.22 | 14.6 | 8.64 | 12.8 | 14.2 | 15.8 | 6.22 |
|  | SC4 | 15.1 | 16.5 | 18.1 | 7.09 | 16.5 | 9.78 | 15.1 | 16.5 | 18.1 | 7.09 |
|  | SC5 | 17.0 | 18.4 | 20.0 | 7.76 | 18.4 | 10.7 | 17.0 | 18.4 | 20.0 | 7.76 |
|  | SC6 | 18.8 | 19.9 | 22.0 | 8.35 | 19.9 | 11.5 | 18.8 | 19.9 | 22.0 | 8.35 |
| | Mean±<br>SD | 6.66±2.48 | 9.83±3.15 | 12.0±3.77 | 13.8±4.37 | 15.4±4.93 | 16.7±5.48 | 17.9±5.94 | 18.9±6.33 | 19.7±6.69 | 20.6±6.96 |
|  |  | P1 vs P2 | P2 vs P3 | P3 vs P4 | P4 vs P5 | P5 vs P6 | P6 vs P7 | P7 vs P8 | P8 vs P9 | P9 vs P10 | - |
| $F_{Packages\_loss}$<br>(%) | SC1 | 3.72 | 4.42 | 3.97 | 1.65 | 2.72 | 2.56 | 3.72 | 4.42 | 3.97 | - |
|  | SC2 | 2.65 | 3.12 | 2.65 | 1.17 | 2.14 | 1.51 | 2.65 | 3.12 | 2.65 | - |
|  | SC3 | 2.28 | 2.29 | 2.26 | 0.86 | 1.92 | 1.14 | 2.28 | 2.29 | 2.26 | - |
|  | SC4 | 1.86 | 1.90 | 1.95 | 0.67 | 1.86 | 0.93 | 1.86 | 1.90 | 1.95 | - |
|  | SC5 | 1.80 | 1.55 | 1.96 | 0.59 | 1.59 | 0.76 | 1.80 | 1.55 | 1.96 | - |
|  | SC6 | 1.52 | 1.31 | 1.66 | 0.46 | 1.32 | 0.70 | 1.52 | 1.31 | 1.66 | - |
| | Mean±<br>SD | 3.17±1.04 | 2.21±0.75 | 1.79±0.63 | 1.53±0.57 | 1.37±0.56 | 1.16±0.48 | 0.96±0.39 | 0.89±0.37 | 038±0.28 | - |

## Notes

### Competing Interest Statement

The authors have declared no competing interest.

